# Neotelomere formation by human telomerase

**DOI:** 10.1101/2022.10.31.514589

**Authors:** Charles G. Kinzig, George Zakusilo, Kaori K. Takai, Titia de Lange

## Abstract

The maintenance of genome integrity requires that telomerase action be limited to telomeres and not convert DSBs into neotelomeres. Using the breakpoint sequence from an apparent germline neotelomere formation event, we developed an assay to detect and quantify telomeric repeat addition at Cas9-programmed DSBs in human cells. The data show that telomerase can add telomeric repeats to DSBs and that this process can generate functional neotelomeres. Neotelomere formation is increased when telomerase is overexpressed, suggesting that in most human cells, low (or absent) telomerase activity limits the deleterious effects of de novo telomere addition. We show that neotelomere formation at DSBs is inhibited by long-range resection and the accompanying activation of ATR signaling. Our findings reveal that telomerase can cause genome instability by generating neotelomeres at DSBs. We propose that neotelomere formation can promote tumorigenesis by ending detrimental breakage-fusion-bridge cycles in cancer cells whose genome alterations engender dicentric chromosomes.

## Introduction

Telomeres define and protect the ends of linear chromosomes. Stable genome maintenance requires that telomeres evade recognition as DNA double-strand breaks (DSBs), which is accomplished by the shelterin complex bound to double-stranded (ds) telomeric repeats (de Lange, 2018). Telomerase counteracts the shortening of telomeres that occurs with cell division due to incomplete DNA replication and nucleolytic resection (Greider and Blackburn, 1985; Hockemeyer and Collins, 2015; Wu et al., 2017). Human telomerase is a ribonucleoprotein complex whose reverse transcriptase component (TERT; (Lingner et al., 1997)) uses its bound RNA component (hTR; (Feng et al., 1995)) to add TTAGGG repeats to the telomeric single-stranded (ss) 3′ overhang (reviewed in (Wu et al., 2017). The template region of hTR contains 11 nucleotides complementary to the telomeric TTAGGG repeats (Feng et al., 1995), which enable telomerase to anneal to the 3′ end of the telomere and serve as a template for repeat addition (Ghanim et al., 2021; Liu et al., 2022; Nguyen et al., 2018; Sekne et al., 2022).

Just as telomeres must not be recognized as DNA damage, telomerase must not convert DSBs into telomeres. If telomerase initiates de novo telomere synthesis at a DSB and successfully creates a functional telomere, the genomic material distal to the break—and thus no longer physically linked to the centromere—will be lost. A priori, three factors could restrict human telomerase to telomeres: the base-pairing of hTR with the 3′ telomeric overhang, the low level or absence of telomerase in most cells, and the requirement for telomerase to be recruited to telomeres by TPP1 (Hockemeyer and Collins, 2015; Zhong et al., 2012; Nandakumar et al., 2012; Abreu et al., 2010).

However, in some organisms, telomerase activity at non-telomeric sites is a physiologic function of the enzyme. For example, in the ciliate *Tetrahymena thermophila*, telomerase stabilizes the DNA fragments formed by programmed cleavage of the micronuclear genome during the generation of the vegetative macronucleus (Fan and Yao, 2000; Fan and Yao, 1996; Yao et al., 1990).

The first hint that telomerase can act at DSBs came from Barbara McClintock’s work on dicentric chromosomes in maize (McClintock, 1938; McClintock, 1941), which are unstable and undergo repeated breakage-fusion-bridge (BFB) cycles in all somatic cells until the broken chromosomes gain functional telomeres in the embryo (McClintock, 1941). Writing to Elizabeth Blackburn in 1983 (McClintock, 1983), McClintock described a maize mutant in which this healing never occurred, suggesting that “this mutant affects the production or the action of an enzyme required for formation of new telomeres.” This presumed enzyme, telomerase, would be identified two years later in extracts from mating *Tetrahymena* (Greider and Blackburn, 1985).

In the budding yeast *Saccharomyces cerevisiae*, the threat to genome integrity posed by telomerase has been studied extensively (Ribeyre and Shore, 2013). The helicase Pif1 prevents inappropriate telomerase activity at budding yeast DSBs, presumably by physically unwinding the telomerase RNA–DNA duplex (Boulé et al., 2005; Boulé and Zakian, 2007; Myung et al., 2001; Schulz and Zakian, 1994). Mec1 also inhibits telomerase at DSBs by phosphorylating Cdc13, a telomeric ssDNA-binding protein that recruits telomerase to telomeres (Zhang and Durocher, 2010).

Sequence analysis of terminal chromosome deletions in patients suggests that telomerase may also threaten DSBs in human cells. In a patient with α-thalassemia due to a truncation of chromosome 16p, telomeric sequences were added directly to a non-telomeric breakpoint sequence, suggesting that telomerase-mediated neotelomere formation may have stabilized the terminal deletion (Wilkie et al., 1990). In vitro telomerase assays demonstrated that the 25-nt sequence preceding the breakpoint (denoted TS) is an excellent telomerase primer (Morin, 1991). Several other cases of terminal chromosome deletions have been reported in which the juxtaposition of breakpoint sequence and telomeric repeats is consistent with neotelomere formation (Ballif et al., 2003; Bonaglia et al., 2011; Flint et al., 1994; Guilherme et al., 2015; Hannes et al., 2010; Horsley et al., 2001; Lamb et al., 1993; Varley et al., 2000; Wong et al., 1997; Yatsenko et al., 2009). Nonetheless, direct evidence that human telomerase can form neotelomeres at DSBs is lacking.

Here, we aimed to determine whether telomerase can add telomeric DNA to DSBs in human cells and how this process is regulated. The data indicate that telomerase can create a functional neotelomere at a Cas9-induced DSB that bears the TS sequence at one 3′ end. The frequency of neotelomere formation is increased upon overexpression of telomerase, suggesting that the low level of telomerase in most human cells minimizes these deleterious events. In addition, neotelomere formation is inhibited by ATR signaling at resected DSBs. We discuss these findings in the context of genome instability during tumorigenesis, where neotelomere formation by telomerase might end BFB cycles as originally proposed by McClintock.

## Results

### Telomerase-mediated TTAGGG repeat addition at Cas9-induced DSBs

To develop a system to measure TTAGGG repeat addition at DSBs, we selected the TS sequence among sites of presumed germline neotelomere formation (Figure S1) because TS is the only breakpoint sequence known to serve as an in vitro primer for telomerase (Morin, 1991). A lentiviral vector (Lenti-sgTS-TaqMan-TS) was constructed that expresses an sgRNA targeting TS (sgTS) and bears the TS sequence flanked by a PAM site such that Cas9 would create a DSB with the TS sequence at the 3’ end (Figure 1A; Figure S2A). An analogous vector expressing an sgRNA targeting luciferase (sgLuc) instead of sgTS served as a negative control. To detect neotelomere formation at the TS site after Cas9 cleavage, a reverse primer was designed to anneal to both the 3’ end of TS and the added TTAGGG repeats assuming the repeat addition occurred in the frame observed in the α-thalassemia patient (Figure 1A; Figure S1; Figure S2A). A TaqMan probe that anneals between the forward primer and TS enabled quantification of neotelomere formation events by qPCR. The assay is not designed to detect functional neotelomeres—merely TTAGGG repeat addition at the programmed DSBs.

**Figure 1.**
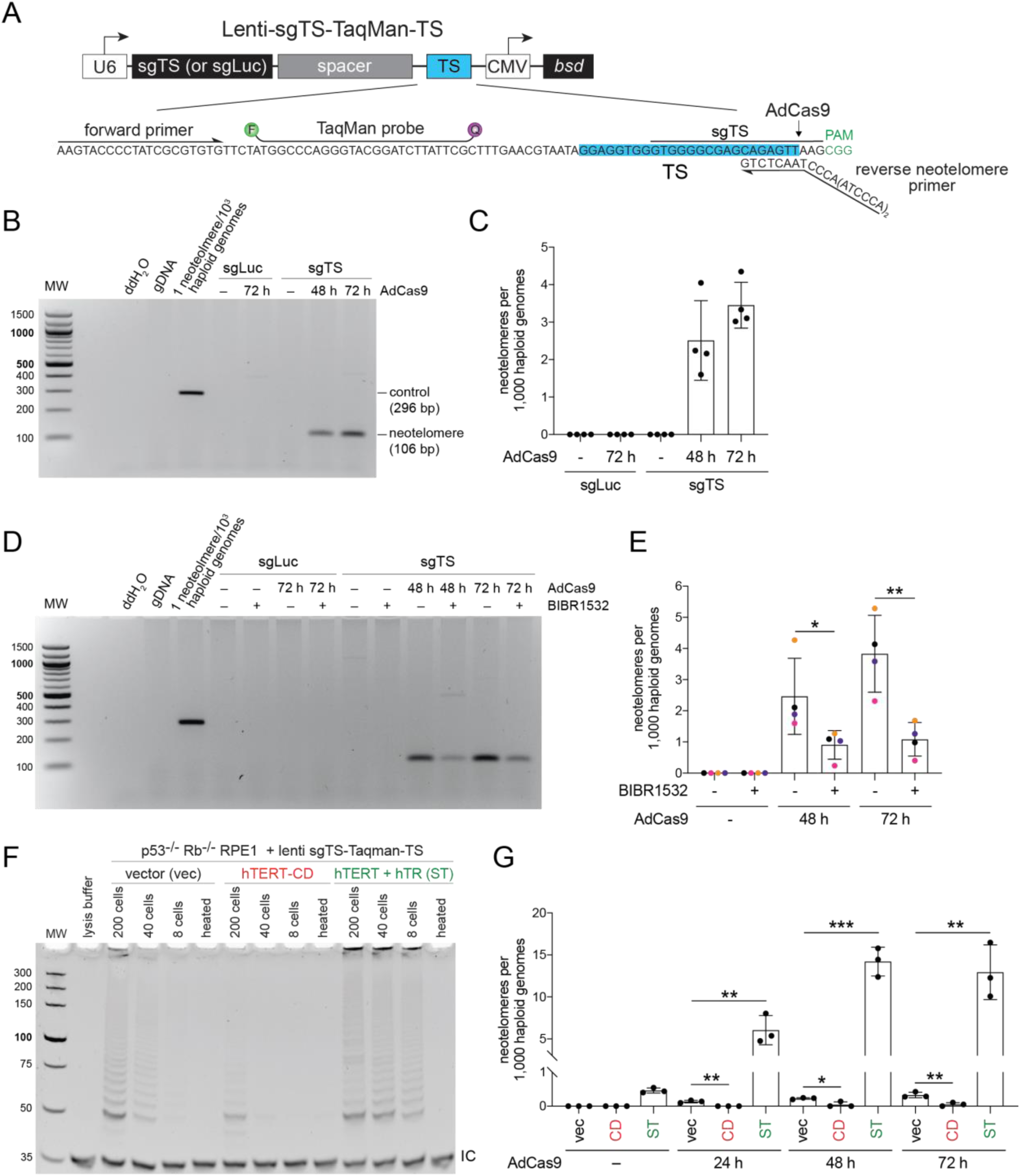
Telomerase-mediated telomeric repeat addition at a Cas9-induced DSB (A) Schematic of the pLenti-sgTS-TaqMan-TS lentiviral construct. U6, human U6 promoter; sgLuc or sgTS, single-guide RNA (sgRNA) cassette; TS, PCR cassette containing the truncation sequence (TS); CMV, cytomegalovirus promoter; *bsd*, blasticidin S deaminase gene. The inset shows the sequence of TS PCR cassette, with the primer and TaqMan binding sites and patient-derived TS sequence indicated. The predicted site of cleavage by Cas9 (AdCas9) within the sequence recognized by the sgRNA is denoted, 3 nucleotides from the protospacer adjacent motif (PAM). (B) Ethidium-bromide-stained agarose gel showing products obtained by endpoint PCR with genomic DNA (gDNA) from super-telomerase HeLa (HeLa-ST) cells expressing targeting (sgTS) or non-targeting (sgLuc) sgRNAs. Genomic DNA was harvested at the indicated times after infection with AdCas9. Sterile water and human genomic DNA (gDNA) were used as negative controls. A plasmid template simulating neotelomere addition in the expected frame of addition was spiked into human gDNA as a positive control. The positions of control (296 bp) and cell-generated (106 bp) neotelomere PCR products are denoted. (C) TaqMan-qPCR quantification of neotelomere formation in (B). (D) Ethidium-bromide-stained agarose gel showing products obtained by endpoint PCR with gDNA from HeLa-ST cells treated with BIBR1532 (20 µM) or DMSO (vehicle). Sample labeling and controls are as in (B). (E) Quantification of neotelomeres in (D) by TaqMan-qPCR. Data points bearing the same color belong to the same biological replicate. (F) Telomeric Repeat Amplification Protocol (TRAP) assay products obtained with extracts from the indicated cells infected with a retroviral vector (vec), a retrovirus expressing catalytically dead (CD) hTERT, and a retrovirus expressing wild-type hTERT plus hTR (ST). TRAP products were resolved by polyacrylamide gel electrophoresis and stained with ethidium bromide. As a control, extracts were heat-inactivated (heat). The position of the internal control (IC) band is noted. (G) Quantification of neotelomeres formed in the RPE1 cells shown in (F) at the indicated times after infection with AdCas9. Mean ± SD of at least 3 biological replicates. * *p* < 0.05, ** *p* < 0.01, *** *p* < 0.001, two-tailed ratio-paired *t*-test in (E) and two-tailed unpaired *t*-test in (G).

Based on qPCR, we estimate that lentivirus-infected HeLa cells contain ∼1.7 copies of the TS cassette per haploid genome (Figure S2B). After infection with Cas9 adenovirus (AdCas9), ∼50% of the TS sites became sensitive to T7 endonuclease I, indicating efficient Cas9 cleavage (Figure S2C). In a mock experiment with a positive control plasmid spiked into genomic DNA, the TaqMan qPCR assay was linear over a range of 10^−4^ to 10 neotelomere formation events per haploid genome (Figure S2D).

We first tested whether the Cas9 cut at TS could lead to neotelomere formation using HeLa cells expressing high levels of telomerase (“super-telomerase” cells, HeLa-ST), which were generated by overexpressing both hTR and hTERT (Cristofari and Lingner, 2006). In these cells, neotelomere products were detected by PCR at 48 and 72 h after AdCas9 infection, whereas no products were detected in uninfected cells or control cells that did not express the sgRNA to TS (Figure 1B). TaqMan qPCR indicated that the cells contained 3–4 neotelomeres per 1,000 haploid genomes at 72 h after AdCas9 infection (Figure 1C).

Treatment of HeLa-ST cells with the telomerase inhibitor BIBR1532 (Pascolo et al., 2002) reduced neotelomere formation 3–4-fold at 72 h (Figure 1D, E), suggesting that the neotelomere products were generated by telomerase. To further test whether the products were due to telomerase activity, we introduced the sgTS-TaqMan-TS lentivirus into p53^-/-^ Rb^-/-^ RPE1 cells (Yang et al., 2017), which have a moderate level of telomerase due to retroviral hTERT expression (Bodnar et al., 1998). These cells were then infected with one of three retroviruses: the empty vector (vec), one encoding a catalytically dead copy of hTERT (hTERT-CD; D712A/V713I (Hahn et al., 1999)), and a vector encoding both wild-type hTERT and hTR to confer super-telomerase activity (Wong and Collins, 2006). Telomeric Repeat Amplification Protocol (TRAP) assays for telomerase activity in extracts of the infected cells showed that hTERT-CD reduced the telomerase activity, whereas overexpression of hTERT and hTR increased it (Figure 1F). Cells infected with the empty vector showed a low level of neotelomere formation (∼0.3 neotelomeres/1,000 haploid genomes at 72 h), which was reduced by expression of hTERT-CD (Figure 1F). The frequency of neotelomere formation at 48 and 72 h after introduction of Cas9 was increased ∼40-fold when telomerase was overexpressed (Figure 1G). The results indicate that the neotelomere products are generated by telomerase and suggest that telomerase levels are limiting for the neotelomere formation events detected by our assay.

#### Evidence for CST-mediated C-strand synthesis during neotelomere formation

To determine whether the TTAGGG repeats synthesized by telomerase at DSBs are converted into a duplex product or remain largely single-stranded, we examined whether the detection of neotelomere formation products was affected by treatment of the genomic DNA with the *E. coli* 3’ exonuclease *Exo*I. As a control for the removal of 3’ ssDNA by *Exo*I, we assessed the genomic DNA samples for the loss of the physiologic telomeric 3’ overhang. As expected, the G-rich ss overhang signal of the bulk telomeres was almost completely removed by *Exo*I, whereas double-stranded telomeric DNA largely resisted degradation (Figure 2A). In the same DNA samples, *Exo*I treatment did not affect the abundance of the detected neotelomere products (Figure 2B), suggesting that the TS-neotelomere junction is double-stranded at the time the genomic DNA is harvested.

**Figure 2.**
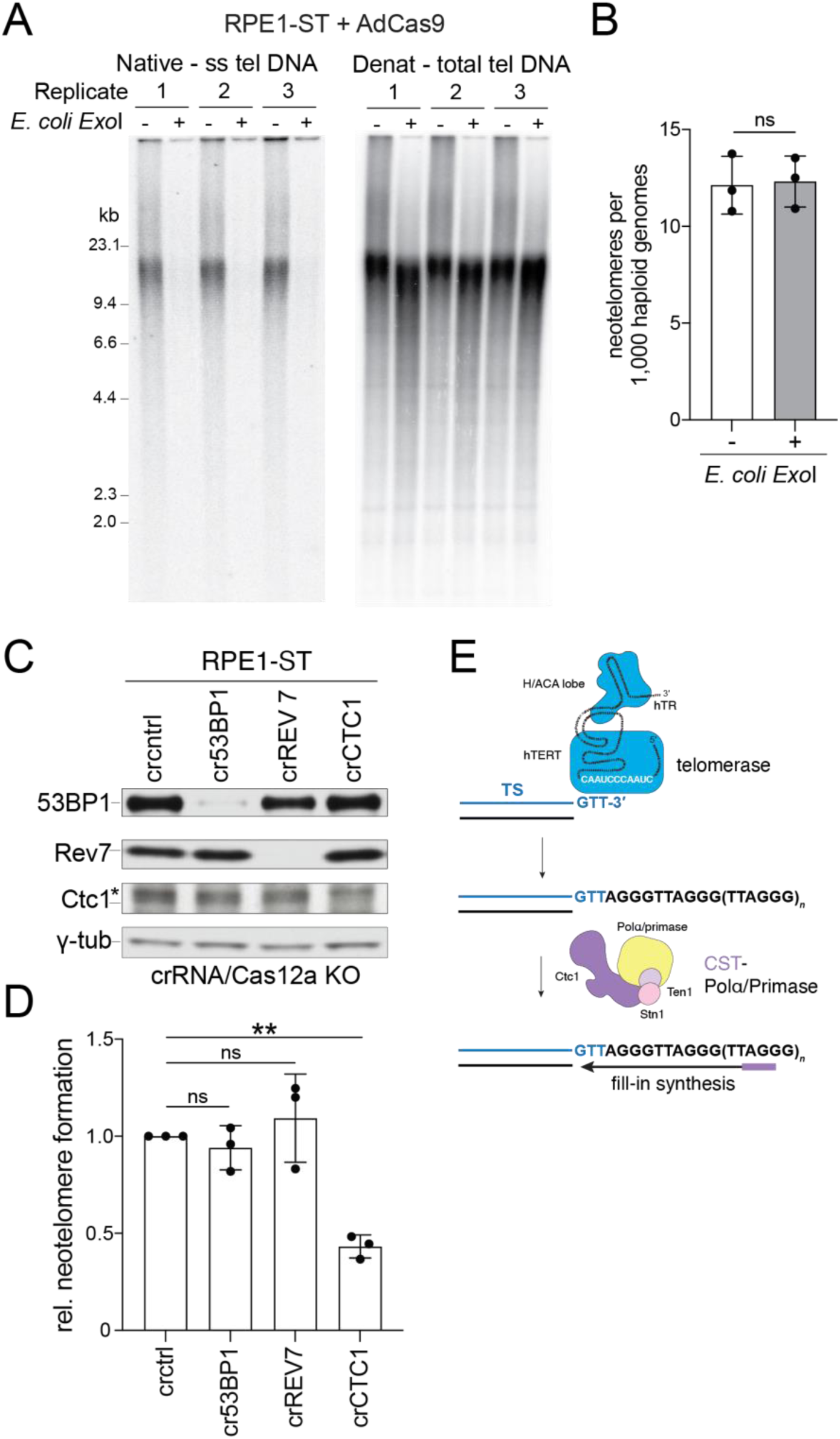
CST-mediated C-strand synthesis during neotelomere formation (A) Telomeric overhang assay showing the effect of *E. coli* exonuclease I (*Exo*I) digestion on three genomic DNA samples from RPE1-ST cells at 48 h after infection with AdCas9. In-gel hybridization was conducted with a radiolabeled C-strand telomeric oligonucleotide probe to detect single-stranded telomeric G-rich overhangs under native conditions (left) and total telomeric DNA after in situ denaturation of the DNA and re-probing with the same oligonucleotide (right). (B) Quantification of neotelomeres by TaqMan qPCR of genomic DNA samples from (A) with or without *Exo*I treatment. (C) Western blot for the indicated proteins performed on whole-cell lysates from RPE1-ST cells treated with Cas12a and CRISPR RNAs (crRNAs) targeting 53BP1, REV7, or CTC1 or a non-targeting control (cntrl) crRNA. γ-tub, loading control; * non-specific band. (A) (D) Quantification of neotelomere formation by TaqMan qPCR in the RPE1-ST cells shown in (C) normalized to cells treated with the control crRNA. (B) (E) Model for G-strand synthesis by telomerase (top) and C-strand synthesis by CST-Polα/primase (bottom) at the TS site. Mean ± SD of 3 biological replicates. ns *p* > 0.05, ** *p* < 0.01, two-tailed ratio-paired *t*-test in (B) and (D).

At telomeres, the proximal part of the ss TTAGGG repeats synthesized by telomerase is rendered double-stranded through post-replicative synthesis of the C-rich telomeric repeat strand (reviewed in (de Lange, 2018)). This fill-in synthesis takes place several hours after telomerase has extended the G-rich strand (Zhao et al., 2009) and is mediated by Ctc1-Stn1-Ten1 (CST), presumably through recruitment of its associated Polα/primase (Wang et al., 2012; Wu et al., 2012). Similarly, CST-Polα/primase can perform fill-in synthesis at resected DSBs, where it is recruited by the 53BP1-Rif1-Shieldin cascade ((Mirman et al., 2018); reviewed in (Mirman and de Lange, 2020; Mirman et al., 2022)).

To test whether CST-Polα/primase fill-in affects the detection of neotelomere formation, we used an orthogonal CRISPR/Cas12a system from *Acidaminococcus* (Zetsche et al., 2015; Zetsche et al., 2017) to knockout (KO) CTC1. The targeting was done in Lenti-sgTS-TaqMan-TS-infected p53^-/-^ Rb^-/-^ RPE1 cells overexpressing hTERT and hTR (Figure 1F), hereafter referred to as RPE1-ST cells. CTC1 KO caused a two-fold reduction in neotelomere formation (Figure 2C, D). This result is consistent with CST-Polα/primase fill-in converting the product of telomerase into duplex DNA, which is presumably more stable since it would enable recruitment of the protective shelterin complex (Figure 2E).

Interestingly, bulk KO of 53BP1 or the Rev7 subunit of shieldin did not affect neotelomere formation (Figure 2C, D), suggesting that CST-Polα/primase can act at TTAGGG repeats independent of this mode of recruitment. Similarly, a recent report noted 53BP1-independent Polα fill-in activity at DSBs (Paiano et al., 2021). The results also argue against the possibility that CST, which is an inhibitor of telomerase at telomeres (Chen et al., 2012), blocks telomerase at DSBs. However, it is possible that CRISPR targeting of CTC1 had a dual effect, increasing neotelomere formation by releasing telomerase from CST-mediated inhibition while simultaneously diminishing the stability of the telomerase product due to a lack of fill-in synthesis.

#### Telomerase mediates a terminal truncation and functional neotelomere formation

To determine whether the observed addition of TTAGGG repeats to the Cas9-induced DSBs can create functional neotelomeres, we used homologous recombination at CRISPR/AsCpf1 cleavage sites to insert a TS cassette construct at the *LUC7L* locus on the distal part of chromosome 16p of HeLa-ST cells (Figure 3A). The Cas9-cleavable TS sequence is positioned centromeric to HSV thymidine kinase (TK) so that neotelomere formation at TS would lead to loss of TK and render cells resistant to ganciclovir. A single HeLa-ST clone with the correct insertion was identified (clone 48; Figure S3A, B) and infected with an sgTS-expressing lentivirus. AdCas9 was delivered to the cells, and ganciclovir-resistant clones were isolated. The clones were screened by PCR for telomeric repeat addition at TS, and some clones were further analyzed for retention of the centromeric portion of the TS cassette and loss of the telomeric portion (Figure S3C, D). Of the 67 ganciclovir-resistant clones analyzed, 21 (31%) showed telomeric repeat addition at TS in endpoint PCR (Figure S3D).

**Figure 3.**
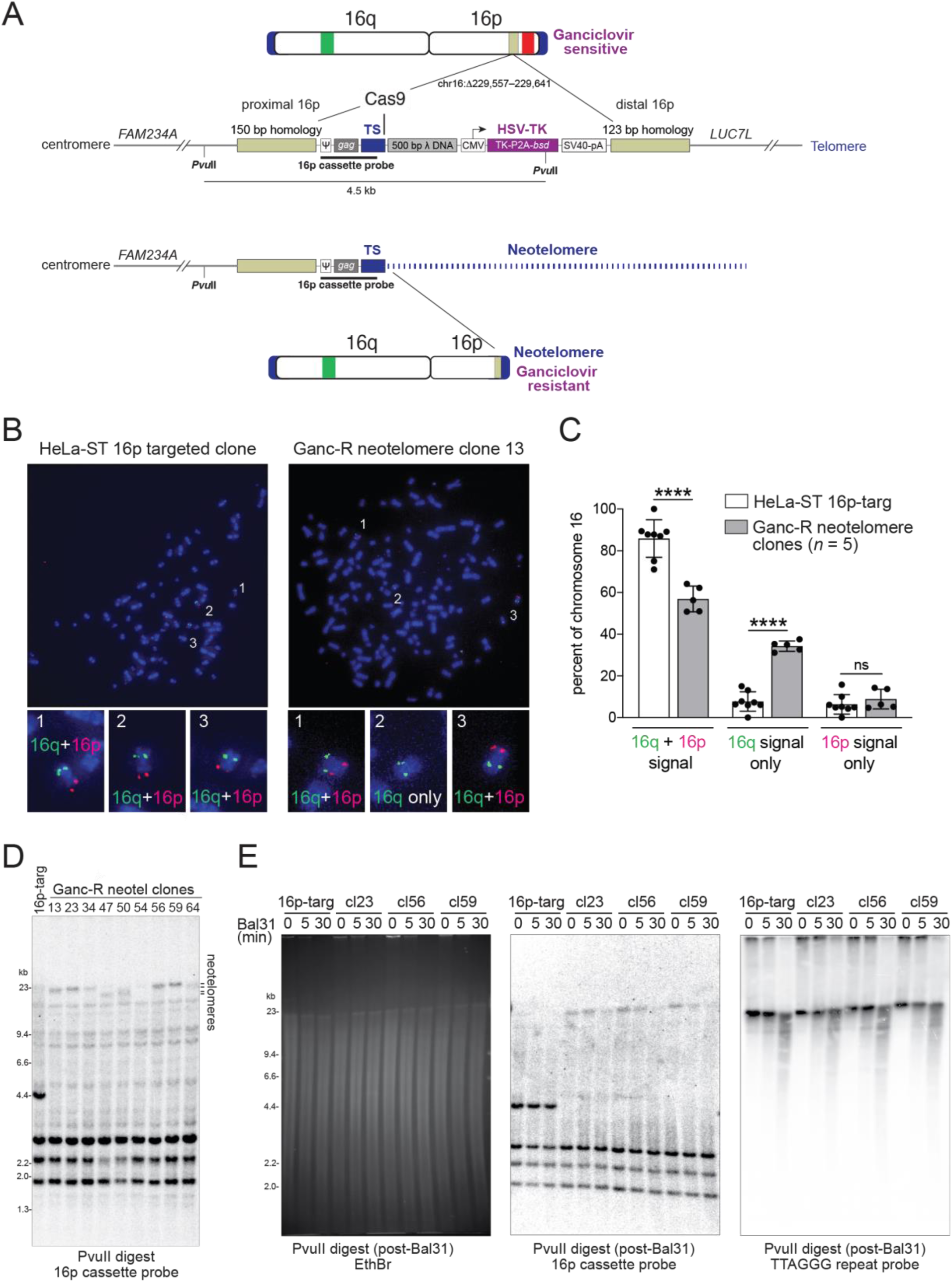
Functional neotelomere formation in HeLa-ST cells (A) Experimental strategy to select for cells in which telomerase has formed a functional telomere at a DSB. A schematic of the CRISPR/Cas12a-edited chromosome 16 in the HeLa-ST 16p targeted clone (16p-targ) is shown, indicating the position of the knock-in cassette (tan) between *FAM234A* and *LUC7L* and the approximate locations of the 16q (green) and 16p (red) FISH probes. The knock-in cassette contains two homology arms and a Cas9-cleavable TS site separated by 500 bp bacteriophage λ DNA from herpes simplex virus (HSV) thymidine kinase (TK) gene, which sensitizes cells to ganciclovir (Ganc). Positions of *Pvu*II sites and a 904-bp Southern blotting probe are indicated. Functional neotelomere formation at TS is predicted to result in loss of TK expression, ganciclovir resistance, loss of the 16p FISH signal, and a change in the size of the PvuII restriction fragment. ψ, MMLV packaging signal; CMV, cytomegalovirus promoter; P2A, 2A peptide from porcine teschovirus-1; *bsd*, blasticidin S deaminase gene; SV40-pA, simian virus 40 polyadenylation signal. The inset shows the sequence of the TS PCR cassette, with the primer and TaqMan binding sites and patient-derived TS sequence indicated. (A) (B) Representative examples of metaphase FISH (see (A)) performed on the parental HeLa-ST 16p targeted clone (16p-targ) bearing the uncleaved TS site and one of the ganciclovir-resistant (Ganc-R) daughter clones (clone 13) identified as potentially harboring a neotelomere based on PCR. The three copies of chromosome 16 are numbered and shown at higher magnification below. (B) Quantification of metaphase FISH in (B). Copies of chromosome 16 were identified by hybridization with either the p or q arm probe and scored as hybridizing with both the p and q arm probes, only the p arm probe, or only the q arm probe. Data plotted are mean ± SD of eight technical replicates for the HeLa-ST 16p targeted clone (23 metaphases per replicate) and means ± SD for pooled data derived from five Ganc-R daughter clones: clone 13, 3 replicates with 18 metaphases each; clone 34, 2 replicates with 20 metaphases each; clone 50, one replicate with 27 metaphases; clone 54, one replicate with 20 metaphases; and clone 64, one replicate with 25 metaphases. ns *p* > 0.05, **** *p* < 0.0001, two-tailed unpaired *t*-test. (C) Southern blot performed on *Pvu*II-digested genomic DNA isolated from the parental HeLa-ST 16p targeted clone (16p-targ) and nine Ganc-R neotelomere clones. The membrane was hybridized with the Klenow-labeled radioactive 16p cassette probe indicated in (A). The position of the neotelomeres is indicated. (D) Southern blot of genomic DNA of the indicated clones treated with Bal31 exonuclease for the indicated times, followed by digestion with *Pvu*II. Left: Ethidium-bromide-stained gel; middle: blot hybridized as in (D); right: same blot reprobed with a radiolabeled telomeric oligonucleotide.

Metaphase spreads from six clones with PCR evidence of neotelomere formation were analyzed by fluorescence in situ hybridization (FISH) to determine whether there was loss of the distal segment of chromosome 16p (Figure 3A–C; Figure S3E). FISH with probes to the q arm and a subtelomeric segment of the p arm (Figure 3A) showed that the parental cell line had three intact copies of chromosome 16 (Figure 3B). By contrast, five of the six neotelomere clones (clones 13, 34, 50, 54, and 64) showed loss of the distal p arm of one of the three copies of chromosome 16 (Figure 3B, C). In these neotelomere clones, the percentage of copies of chromosome 16 staining for both the q and p FISH probes decreased, whereas the percentage of chromosomes staining for only 16p increased (Figure 3C). The metaphase spreads of a sixth neotelomere clone (clone 23) showed one wild-type chromosome 16 and a derivative chromosome 16 with only the 16q signal, consistent with neotelomere formation (Figure S3E). However, this clone was unusual since the signal for the subtelomeric segments of 16p appeared to be attached to another chromosome (Figure S3E).

As an orthogonal approach to determine whether the clones carried new telomeres at the TS site, Southern blots of *Pvu*II-digested genomic DNA were probed with a 904-bp radiolabeled fragment that should detect a 4.5-kb fragment if the TS cassette is intact and a larger fragment if a telomere is added at TS (Figure 3A). Each of the 9 clones analyzed showed a large fragment (∼20 kb), whereas the parental cells showed the expected 4.5-kb *Pvu*II fragment (Figure 3D). The clones with the large fragments included clone 23, consistent with this clone carrying a chromosome 16 lacking the distal segment of 16p (Figure S3E). The length of the *Pvu*II fragments in these clones is as expected if telomerase in HeLa-ST cells added the reported ∼600 bp/telomere/PD (Cristofari and Lingner, 2006) over the period of ∼45 days between Cas9 introduction and the isolation of genomic DNA.

To determine whether the large fragments corresponded to chromosome ends, the genomic DNA was treated with the Bal31 exonuclease prior to *Pvu*II digestion (de Lange and Borst, 1982). The ∼20 kb fragments were sensitive to Bal31, indicating that they represent terminal fragments (Figure 3E). The neotelomeric *Pvu*II fragments ran at approximately the same position as the bulk telomeres in the clones and showed a similar sensitivity to Bal31 (Figure 3E). These results establish that telomeric repeat addition at TS can result in the formation of functional neotelomeres.

### Regulation of neotelomere formation

To determine whether DSB repair pathways compete with telomerase at the Cas9-cleaved TS site, we interfered with classical non-homologous end joining (c-NHEJ) by targeting Lig4 and Ku70/80 with CRISPR, homology-directed repair (HDR) by targeting BRCA2, alternative end joining (a-EJ) by targeting Lig3 and PARP1, and single-strand annealing (SSA) by targeting Rad52 (Figure 4A). None of these interventions increased neotelomere formation (Figure 4B), suggesting that there is no single DSB repair pathway that represents a major competitor for neotelomere formation.

**Figure 4.**
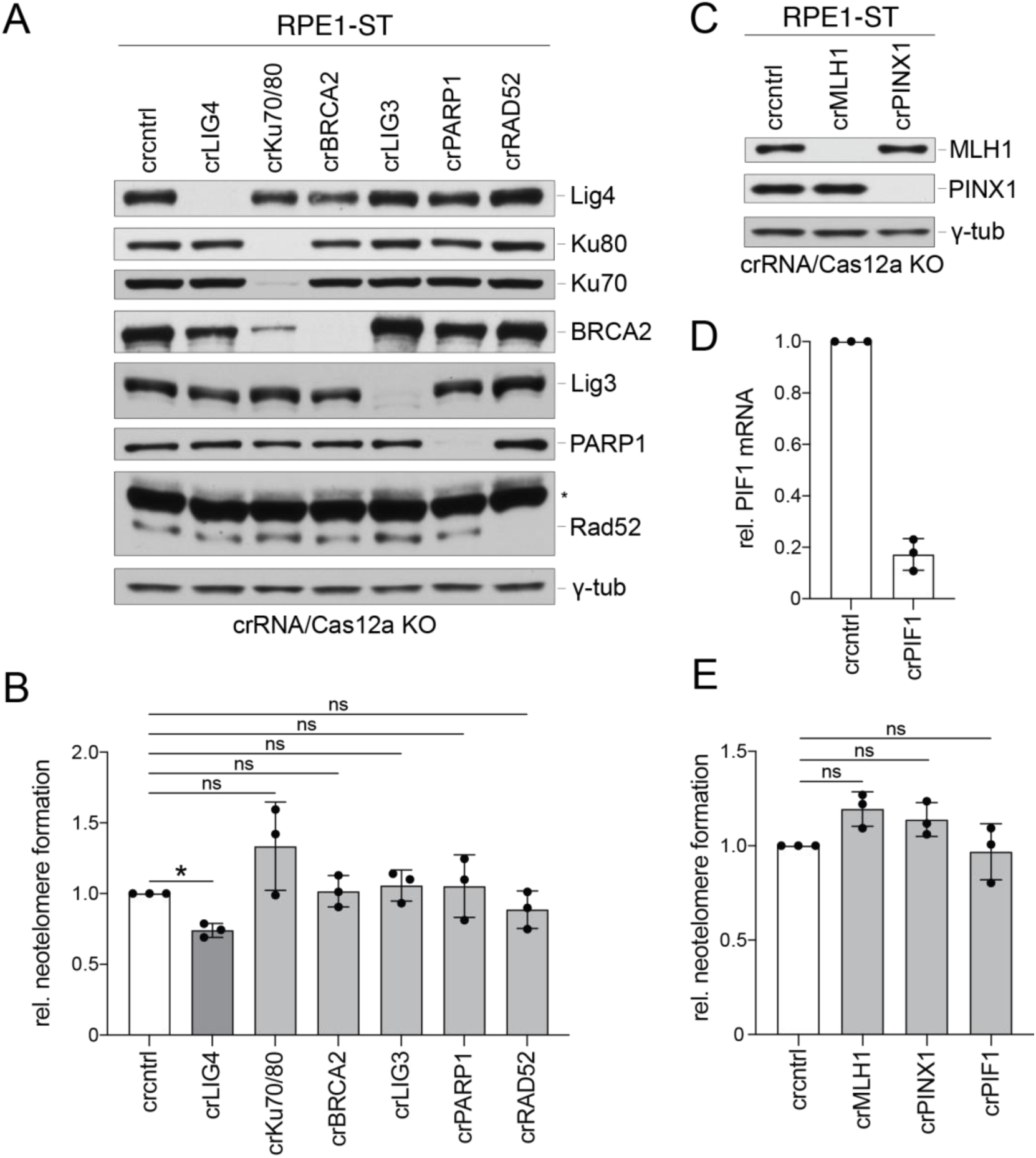
Disruption of individual DSB repair pathways does not affect neotelomere formation (A) Western blot for the indicated proteins performed on whole-cell lysates from RPE1-ST cells treated with Cas12a and CRISPR RNAs (crRNAs) targeting LIG4, Ku70/80 (combination of crKu70 and crKu80), BRCA2, LIG3, PARP1, or RAD52 or a non-targeting control (cntrl) crRNA. γ-tub, loading control; * non-specific band. (B) Quantification of relative neotelomere formation by TaqMan qPCR in the RPE1-ST cells shown in (A) normalized to cells treated with the control crRNA. (C) Western blot for the indicated proteins performed on whole-cell lysates from RPE1-ST cells treated with Cas12a and crRNAs targeting MLH1 or PINX1 or a non-targeting control (cntrl) crRNA. γ-tub, loading control. (D) Abundance of PIF1 mRNA as assessed by RT-qPCR in RPE1-ST cells treated with Cas12a and a crRNA targeting PIF1 or the control crRNA. Relative PIF1 mRNA levels were normalized to GAPDH mRNA. (E) Quantification of neotelomere formation in RPE1-ST cells treated with crRNAs targeting MLH1, PINX1, or PIF1 normalized to that of cells treated with a non-targeting control (cntrl) crRNA. Mean ± SD of 3 biological replicates. ns *p* > 0.05, * *p* < 0.05, two-tailed ratio-paired *t*-test in (B) and (E).

We next tested whether neotelomere formation was affected by MLH1 or PINX1. MLH1 has been proposed to inhibit telomerase-dependent interstitial telomeric repeat insertions (Jia et al., 2017), and PINX1 has been reported to reduce telomerase activity and cause telomere shortening in telomerase-positive cells (Zhou and Lu, 2001). However, CRISPR targeting of neither MLH1 nor PINX1 affected the frequency of neotelomere formation in our assay (Figure 4C–D). Similarly, targeting of PIF1, the ortholog of a major inhibitor of yeast telomerase at DSBs (Myung et al., 2001; Schulz and Zakian, 1994), did not increase neotelomere formation either (Figure 4D–E).

#### Neotelomere formation is repressed by resection, RPA, and ATR signaling

Budding yeast neotelomere formation is increased in strains that lack long-range resection due to mutations in Exo1 and the RecQ helicase Sgs1, the latter of which enables 3’ overhang generation by the DNA2 nuclease (Chung et al., 2010; Lydeard et al., 2010). Therefore, we monitored neotelomere formation after disrupting Exo1 and DNA2, which acts together with BLM and WRN, the mammalian orthologs of Sgs1 (reviewed in (Cejka and Symington, 2021)). Exo1 was targeted with CRISPR/Cas12a and DNA2 was knocked down with an shRNA (Figure 5A). Exo1 loss alone led to a 1.5-fold increase in neotelomere formation, whereas DNA2 knockdown alone led to a 2.7-fold increase (Figure 5B). Combined depletion of Exo1 and DNA2 increased neotelomere formation 3.6-fold, although this effect was not significantly different from that of DNA2 knockdown alone (Figure 5B).

**Figure 5.**
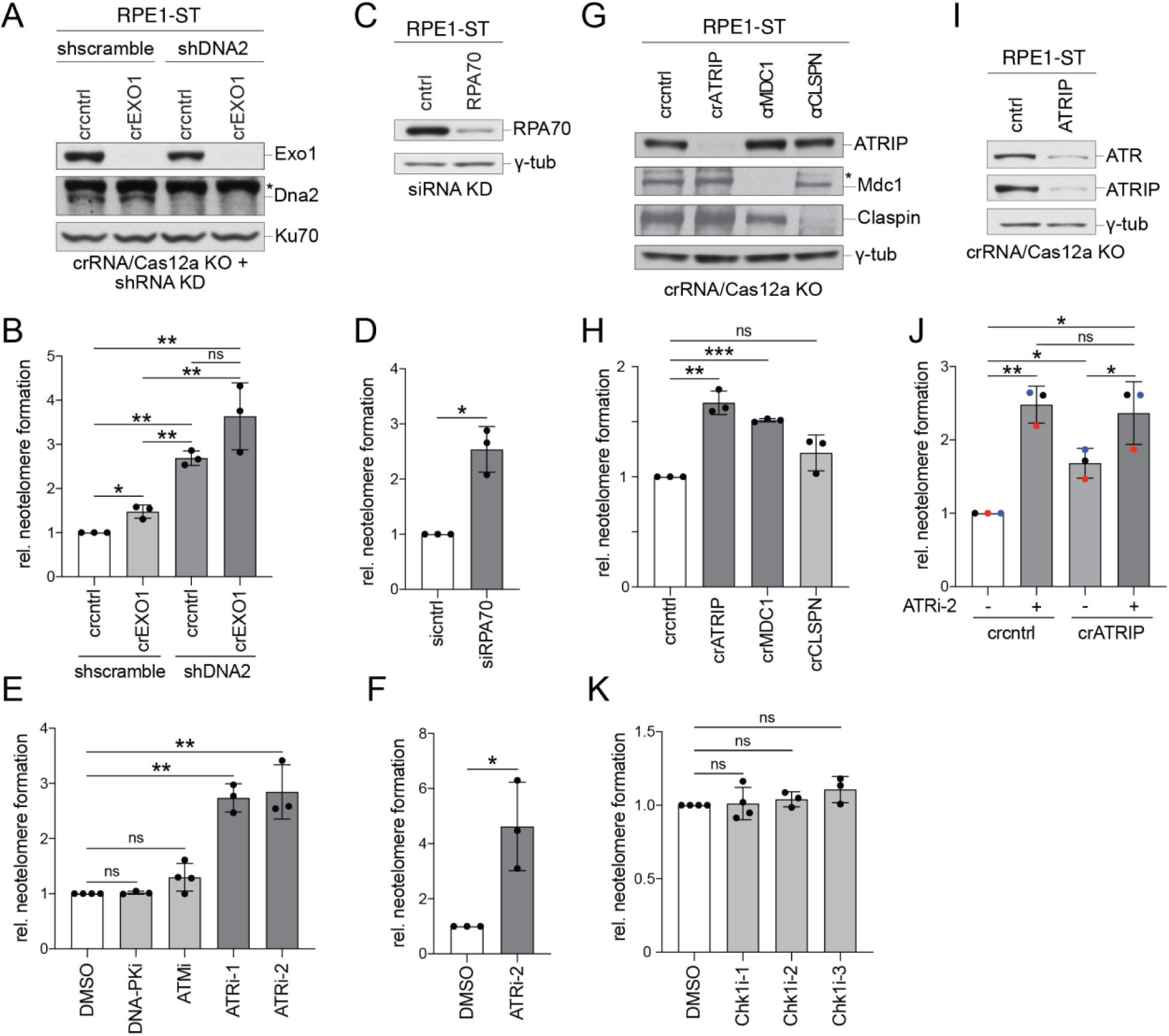
Repression of neotelomere formation by resection, RPA, and ATR signaling (A) Western blot for the indicated proteins performed on whole-cell lysates from RPE1-ST cells treated with Cas12a and CRISPR RNAs (crRNAs) targeting EXO1 or a non-targeting control (cntrl) crRNA followed by shRNAs targeting DNA2 or a scramble shRNA control. Ku70 serves as the loading control; * non-specific band. (B) Quantification of relative neotelomere formation by TaqMan qPCR in cells in (A) normalized to cells treated with the control crRNA and scramble shRNA. (C) Western blot for RPA70 performed on whole-cell lysates from RPE1-ST cells treated with a control (cntrl) siRNA or two siRNAs targeting RPA70. γ-tub, loading control. (D) Quantification of relative neoteolomere formation by TaqMan qPCR in cells in (C) normalized to cells treated with the control siRNA. (E) Quantification of relative neotelomere formation by TaqMan qPCR in RPE1-ST cells treated with DMSO (vehicle) or the indicated inhibitors. Neotelomere frequency was normalized to the DMSO control sample. (F) Quantification of relative neotelomere formation by TaqMan qPCR in RPE1 cells treated with DMSO (vehicle) or ATRi-2. Neotelomere frequency was normalized to the DMSO control sample. (G) Western blot for the indicated proteins performed on whole-cell lysates from RPE1-ST cells treated with Cas12a and crRNAs targeting ATRIP, MDC1, or CLSPN or a control (cntrl) crRNA. γ-tub, loading control. (H) Quantification of relative neotelomere formation by TaqMan qPCR in cells in (G) normalized to cells treated with the control crRNA. (I) Western blot for the indicated proteins performed on whole-cell lysates from RPE1-ST cells treated with Cas12a and crRNAs targeting ATRIP or a control (cntrl) crRNA. γ-tub, loading control. (J) Quantification of relative neotelomere formation by TaqMan qPCR in cells in (I) with DMSO (vehicle) or ATRi-2 normalized to cells treated with the control crRNA and DMSO. Data points bearing the same color belong to the same biological replicate. (K) Quantification of relative neotelomere formation by TaqMan qPCR in RPE1-ST cells treated with DMSO (vehicle) or the indicated Chk1 inhibitors normalized to cells treated with DMSO. Mean ± SD of at least 3 biological replicates. ns *p* > 0.05, * *p* < 0.05, ** *p* < 0.01, *** *p* < 0.001, two-tailed ratio-paired *t*-test in (B), (D), (E), (F), (H), (J), and (K). DNA-PKcsi, AZD-6748, 10 µM; ATMi, KU55933, 10 µM; ATRi-1, VE-821, 10 µM; ATRi-2, Gartisertib/M4344, 0.3 µM in (E) and (F) and 1 µM in (J); Chk1i-1, CHIR124, 0.25 µM; Chk1i-2, MK-8776, 1 µM; Chk1i-3, CCT245737, 1 µM.

Another repressor of neotelomere formation in budding yeast is the ssDNA-binding protein RPA (Chen and Kolodner, 1999). To determine whether RPA suppressed human telomerase at DSBs in our system, we measured the frequency of neotelomere formation in RPE1-ST cells treated with two small interfering RNAs (siRNAs) targeting RPA70 (Figure 5C). RPA70 silencing elicited a 2.5-fold increase in neotelomere formation at the TS site (Figure 5D).

The increase in neotelomere formation by inhibition of resection and RPA suggested that telomerase might be controlled by ATR signaling at resected DSBs (reviewed in (Saldivar et al., 2017)). To determine the role of ATR signaling, neotelomere formation assays were performed in RPE1-ST cells treated with two ATR inhibitors (VE-821 and Gartisertib/M4344). ATR inhibition nearly tripled the frequency of neotelomere formation, whereas inhibitors of DNA-PKcs or ATM had no effect (Figure 5E). In RPE1 cells not endowed with super-telomerase activity, ATR inhibition increased neotelomere formation by more than 4-fold (Figure 5F). Consistent with a role for ATR in suppressing telomerase at DSBs, bulk CRISPR targeting of the obligatory ATR partner, ATRIP, resulted in a modest but significant increase in neotelomere formation (Figure 5G, H). This modest increase is likely due to inefficient removal of ATRIP since ATRi treatment of ATRIP-targeted cells increased the frequency of neotelomere formation to the level observed in control cells treated with ATRi (Figure 5I, J).

The inhibition of neotelomere formation by ATR signaling appeared to not require its downstream effector kinase Chk1 since three Chk1 inhibitors failed to affect the frequency of neotelomere formation (Figure 5K). Targeting of claspin, which mediates the activation of Chk1 by ATR, also had no effect (Figure 5G, H). Therefore, ATR is unlikely to affect neotelomere formation through Chk1. By contrast, the ability of ATR to induce DNA damage foci may be involved in the control of neotelomere formation since bulk CRISPR targeting of a constituent of DNA damage foci, MDC1, increased neotelomere formation (Figure 5G, H).

We queried several potential ATR targets that might affect neotelomere formation, focusing on ssDNA-binding proteins that might block telomerase from accessing the 3′ end, helicases that might remove telomerase, and 3′ flap endonucleases that might remove the telomerase product from the DNA end before it is rendered double-stranded. However, no change in neotelomere formation was observed after bulk CRISPR targeting of the ssDNA-binding proteins hSSB1 and RadX (Figure S4A, B), the RecQ helicases BLM and WRN (Figure S4C, D), and the nuclease scaffold SLX4 and the XPF 3′ flap endonuclease (Figure S4E–G). Loss of the FANCJ helicase or Rad51 recombinase led to modest increases in the frequency of neotelomere formation (Figure S4A-D). Perhaps other targets of ATR inhibit neotelomere formation, or ATR acts through multiple pathways. Further work will be required to determine the details of ATR-mediated interference with neotelomere formation.

### The role of MRN/CtIP in regulating neotelomere formation

Since human telomerase does not act at blunt ends (Lei et al., 2005; Rivera and Blackburn, 2004), it might be expected that neotelomere formation would require the MRN/CtIP-dependent endonucleolytic cleavage reaction that creates a short 3′ overhang at DSBs (reviewed in (Stracker and Petrini, 2011) and (Cejka and Symington, 2021)). On the other hand, this initial resection step is required for the long-range resection that may ultimately lead to ATR-dependent inhibition of neotelomere formation. Interestingly, bulk CRISPR targeting of the Mre11 subunit of MRN/CtIP resulted in a 2-fold increase in neotelomere formation in RPE1-ST cells and also led to an increase in RPE1 cells lacking super-telomerase activity (Figure 6A–D). The increase in neotelomere formation was also observed with bulk CRISPR targeting of CtIP (Figure 6A, B), indicating that the nucleolytic activity of MRN/CtIP is the critical aspect of this multifunctional complex in this context.

**Figure 6.**
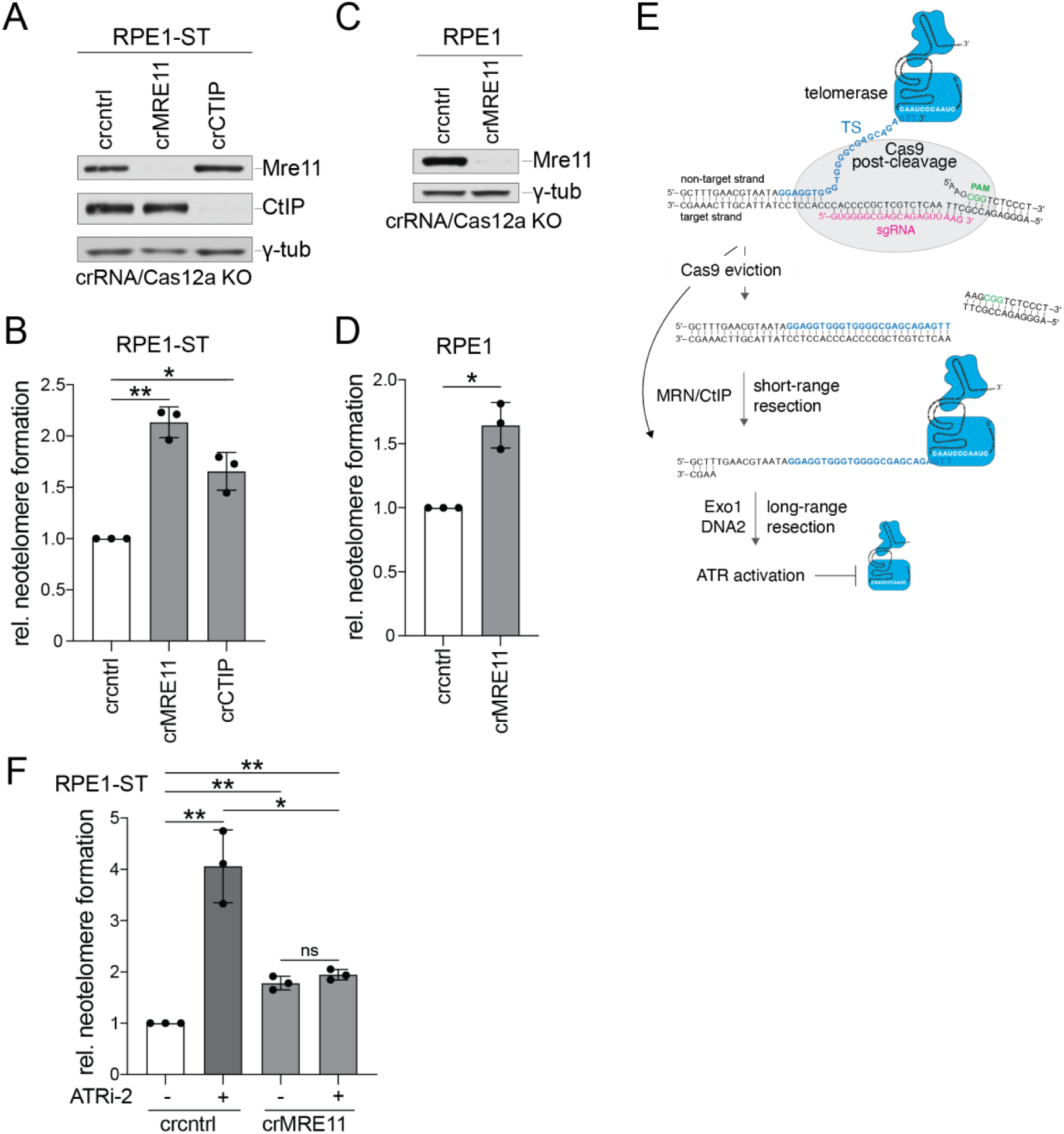
Regulation of neotelomere formation by MRN/CtIP (A) Western blot for the indicated proteins performed on whole-cell lysates from RPE1-ST cells treated with Cas12a and CRISPR RNAs (crRNAs) targeting MRE11 or CtIP or a control (cntrl) crRNA. γ-tub, loading control. (B) Quantification of relative neotelomere formation by TaqMan qPCR in cells in (A) normalized to cells treated with the control crRNA. (C) Western blot for the indicated proteins performed on whole-cell lysates from sgTS-TaqMan-TS RPE1 cells treated with Cas12a and crRNAs targeting MRE11 or a control (cntrl) crRNA. γ-tub, loading control. (D) Quantification of relative neotelomere formation by TaqMan qPCR in cells in (C) normalized to cells treated with the control crRNA. (E) Model for the regulation of neotelomere formation at a Cas9-induced DSB in the presence and absence of MRN/CtIP. The schematic shows the two proposed substrates for telomerase. When Cas9 persists, the extruded non-target strand is the substrate, and MRN/CtIP is not required. When Cas9 is evicted, short-range resection by MRN/CtIP is required to generate the free 3’ end for telomerase priming. Long-range resection can activate ATR, which inhibits neotelomere formation. (F) Quantification of relative neotelomere formation by TaqMan qPCR in sgTS-TaqMan-TS RPE1 cells treated with Cas12a and crRNAs targeting MRE11 or a control (cntrl) crRNA with or without ATR inhibition (ATRi). Neotelomere frequencies were normalized to the sample treated with the control crRNA and DMSO (vehicle). Mean ± SD of 3 biological replicates. ns *p* > 0.05, * *p* < 0.05, ** *p* < 0.01, two-tailed ratio-paired *t*-test in (B), (D), and (F). ATRi, Gartisertib/M4344, 0.3 µM.

The increase in neotelomere formation upon loss of the initial resection step is unexpected given that telomerase requires a ∼5-nt 3′ overhang in vitro (Lei et al., 2005; Rivera and Blackburn, 2004) and suggested that our system provides a substrate for telomerase in the absence of resection. It has been reported that Cas9 in complex with its sgRNA can remain associated with its DNA target after the cleavage reaction (Sternberg et al., 2014). Structural analysis suggests that in the post-cleavage configuration, the strand that is not complementary to the sgRNA protrudes (Zhu et al., 2019). If such a complex is formed at the TS cleavage site, it will expose the telomerase addition site in ss form (Figure 6E). We imagine that this is the substrate that enables neotelomere formation in cells lacking Mre11 or CtIP, where resection initiation is limited. It is possible that MRN/CtIP can remove a persistent Cas9-sgRNA-DNA complex by analogy to the complex’s activity on other bulky protein-DNA adducts (Figure 6E) (Di Virgilio and Gautier, 2005; Hoa et al., 2016; Liao et al., 2016), potentially prolonging the opportunity for telomerase to act on the post-cleavage Cas9 complex when MRN/CtIP is removed.

The hypothetical telomerase substrate in the Cas9 post-cleavage complex (Figure 6E) contains too little single-stranded DNA for ATR activation (MacDougall et al., 2007). Therefore, it is predicted that in Mre11 KO cells where this substrate may be the predominant priming site available to telomerase, ATR would not inhibit neotelomere formation. Consistent with this prediction, treatment of Mre11-deficient cells with an ATR inhibitor had no effect on neotelomere formation (Figure 6F).

The considerable increase in neotelomere formation upon ATRi treatment of MRN/CtIP-proficient cells (Figure 5E, F) suggests that neotelomere formation also takes place at the resected DSB created after Cas9 is removed. Thus, our assay system identifies two types of DNA ends where neotelomere formation can occur: DSBs where Cas9/sgRNA binding has displaced the 3′ end and DSBs evacuated by Cas9 that have been processed by MRN/CtIP-mediated short-range resection (Figure 6E).

## Discussion

We report that human telomerase can add telomeric DNA to a non-telomeric DNA end and can promote the formation of functional neotelomeres at broken chromosome ends (Figure 7A). Neotelomere formation and the accompanying loss of heterozygosity (LOH) due to terminal deletions pose a threat to genome integrity that is likely minimized by the low telomerase activity in most human cells (Figure 7A). In addition, the data indicate that neotelomere formation is inhibited by the activation of ATR signaling at resected DSBs (Figure 7A).

**Figure 7.**
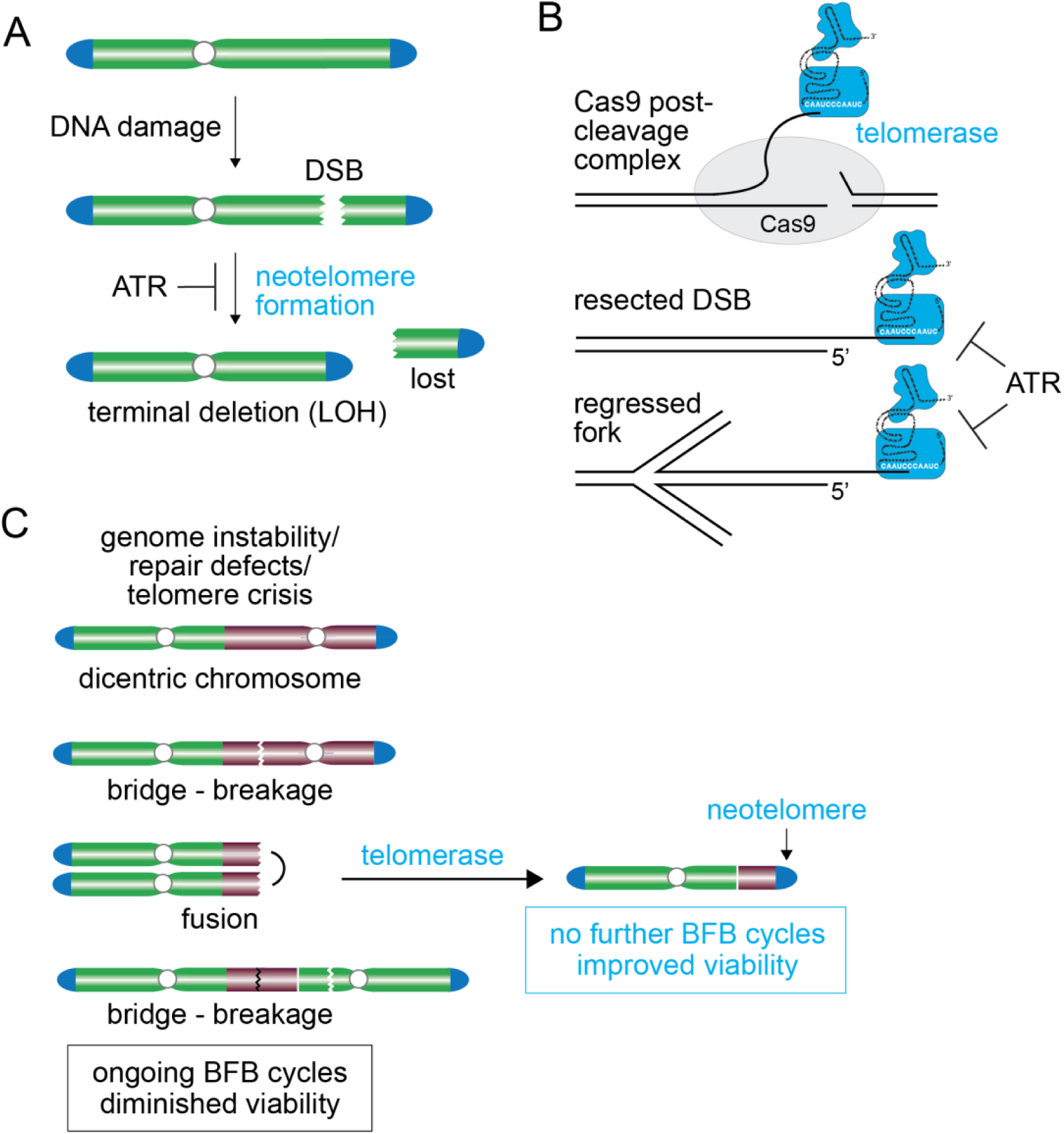
Neotelomere formation by telomerase (A) Model for neotelomere formation at a DSB. The data indicate that ATR signaling represses neotelomere formation at vulnerable DSBs. Neotelomere formation is also limited by the low (or absent) telomerase expression in most human cells. Stable neotelomere formation is predicted to result in loss of the acentric chromosome fragment and a terminal deletion with loss of heterozygosity (LOH) for distal genes. (B) Potential substrates for neotelomere formation. The 3’ end of the extruded non-target strand in the Cas9 post-cleavage complex is proposed to serve as a primer for telomerase. In this setting, ATR is not activated and does not inhibit neotelomere formation. Telomerase can also form neotelomeres at resected DSBs and regressed replication forks, but at these sites, it is inhibited by ATR signaling when resection has created a substrate that activates ATR. (C) Proposed role for neotelomere formation in cancer. Neotelomere formation may end ongoing BFB cycles in genomically unstable cancer clones and thus enhance fitness.

Ultimately, stable neotelomere formation will require sufficient duplex telomeric DNA to be generated for shelterin to bind. If shelterin is not loaded onto the telomeric DNA, it is likely that end-joining reactions will lead to interstitial insertion of the telomerase product. However, once shelterin is associated with the nascent telomere, the telomeric DNA is expected to become protected from the DNA damage response, and the TPP1 subunit of shelterin can recruit telomerase for further extension. The functional neotelomeres observed in our system reached a length similar to the other telomeres in the cells, indicating that they successfully recruited telomerase. The rapid increase in telomere length to many kilobases of telomeric DNA is consistent with prior observations on cells transfected with short telomeric seeds that became elongated to match the length of the host’s telomeres (Hanish et al., 1994).

### Substrates for neotelomere formation

Our assay system for neotelomere formation used a Cas9-induced DSB at a sequence (TS) that was previously suggested to represent a site of neotelomere formation in an α-thalassemia patient. This patient represents one of several potential neotelomere formation events in the human germline. Given the wide variety of sequences where telomerase appears to have acted, it is likely that neotelomere formation is not specific to TS but can take place at many DSBs. Indeed, neotelomere formation was recently observed at an I-SceI cleavage site in mouse embryonic stem cells (Samantha Regan and Maria Jasin, personal communication).

The data argue that our neotelomere formation assay monitors the addition of telomeric repeats at two types of DNA ends (Figure 7B). First, based on the ability of ATR signaling to inhibit neotelomere formation, we deduce that many of the detected events take place at a DNA end that has undergone 5’ resection. However, we also detected neotelomere formation in the absence of MRN/CtIP, the complex that initiates end-resection. We therefore propose that the Cas9-sgRNA-DNA complex, which is thought to persist at DNA ends after cleavage, may also provide a substrate for telomerase (Figure 7B). We imagine that the 3′ end of the non-target strand is accessible to telomerase in this post-cleavage complex. The released non-target strand is likely too short to activate ATR, so telomerase would not be inhibited by ATR signaling in this setting. Accordingly, in the context of therapeutic genome editing by CRISPR-Cas9, CRISPR-Cas12a (Swarts et al., 2017), and prime-editing (Anzalone et al., 2019), it may be prudent to avoid any base-pairing potential between the 3′ end of the non-target strand and the template sequence in hTR. Finally, the ability of telomerase to promote neotelomere formation at DSBs suggests that the enzyme may also act at regressed replication forks (Figure 7B), potentially creating terminal deletions at sites of replication stress.

### Implications for genome instability in cancer

Whole-genome sequencing has revealed numerous structural variations in most cancer genomes (Alexandrov et al., 2020), indicating that most cancers have experienced a multitude of DSBs throughout their development (Gerstung et al., 2020). To what extent such breaks are converted into neotelomeres may depend on the level of telomerase expression during tumorigenesis and the selective advantage afforded by the LOH resulting from terminal deletions. Interestingly, a recent analysis of lung cancer genomes using long-read sequencing methods found evidence for neotelomere formation in ∼20% of the examined cases (Kar-Tong Tan and Matthew Meyerson, pers. comm.).

We speculate that neotelomere formation by telomerase may provide an advantage to incipient cancers that are struggling with ongoing BFB cycles (Figure 7C). Misrepair of DSBs in cancers that are deficient in HDR (e.g., BRCA1/2-deficient cancers) can generate multicentric (radial) chromosomes that can initiate BFB cycles. Interstrand-crosslinks can also generate multicentric chromosomes when their repair is impeded (e.g., in Fanconi anemia (Kottemann and Smogorzewska, 2013)). Similarly, BFB cycles arise during telomere crisis when severely shortened telomeres lack the ability to prevent dicentric formation through telomere-telomere fusions (Cleal et al., 2019; Dewhurst et al., 2021; Maciejowski et al., 2015); reviewed in (Maciejowski and de Lange, 2017)). Most human cancer types show the genomic scars of past BFB cycles, including more than 40 percent of esophageal, gastric, and bladder cancers as well as non-small cell lung cancer and lung squamous cell carcinomas (Hadi et al., 2020). While repeated BFB cycles may accelerate tumor evolution through genome diversification, they may also compromise cellular fitness through loss of essential genes and the detrimental effects of cells undergoing division with dicentric chromosomes. Furthermore, BFB cycles are expected to generate micronuclei containing lagging dicentric chromosomes (or fragments thereof) which will activate cGAS-STING signaling (Ablasser and Chen, 2019; Li and Chen, 2018), promoting recognition and elimination of neoplastic cells by the immune system (Motwani et al., 2019). We propose that by terminating BFB cycles and restoring end-protection to chromosome breaks in genomically unstable tumors, neotelomere formation by telomerase may enable fledgling cancer cells to cope with genome instability.

See also Figures S1 and S2.

See also Figure S3.

See also Figure S4.

## STAR★Methods

Detailed methods are provided in the online version of this paper and include the following:

- Key Resources Table
- Resource Availability

`o` Lead contact
`o` Materials availability
- Experimental Model and Subject Details
- Method Details

`o` Cell culture
`o` Western blotting
`o` Lentiviral and retroviral production and infection
`o` TaqMan-TS-sgTS neotelomere formation assay
`o` Genomic DNA extraction
`o` Endpoint neotelomere PCR reactions
`o` Neotelomere TaqMan qPCR
`o` TRAP assays
`o` *E. coli* exonuclease I digestion
`o` In-gel analysis of single-stranded telomeric DNA
`o` AsCpf1 knockouts
`o` Pif1 RT-qPCR
`o` shRNA knockdown
`o` siRNA knockdown of RPA70
`o` Isolation of HeLa-ST clones containing stable neotelomeres
`o` Metaphase FISH
`o` Southern blotting for the TS cassette
`o` T7 endonuclease I assay
- Quantification and Statistical Analysis

## Supplemental Information

Supplemental information can be found online at *URL*.

## Acknowledgments

HeLa-ST cells were a kind gift from Dr. Joachim Lingner. We thank Dr. John Maciejowski, Dr. Marcin Imieliński, Dr. Dirk Hockemeyer, Dr. David Cortez, Dr. Tom Cech, and members of the de Lange lab for helpful discussion of this work. We recognize the members of the Rockefeller University Genomics Resource Center, especially Bin Zhang, for their assistance with qPCR. This work is supported by grants to T.d.L. from the National Institutes of Health (R35 CA210036), Breast Cancer Research Foundation (BCRF-22-036), and STARR Cancer Consortium (I13-0019). C.G.K. is supported by a F30 Predoctoral Fellowship from the National Cancer Institute of the National Institutes of Health under award number F30CA257419. C.G.K. and G.Z. are supported by a Medical Scientist Training Program grant from the National Institute of General Medical Sciences of the National Institutes of Health under award number T32GM007739 to the Weill Cornell/Rockefeller/Sloan-Kettering Tri-Institutional MD-PhD Program.

## Author Contributions

Conceptualization: C.G.K. and T.d.L; Methodology: C.G.K., G.Z., K.K.T., and T.d.L.; Experiments: C.G.K., G.Z., and K.K.T.; Analysis: C.G.K., G.Z., and K.K.T.; Writing: C.G.K. and T.d.L.

## Supplemental Information

### Key Resources Table

#### Cell Lines

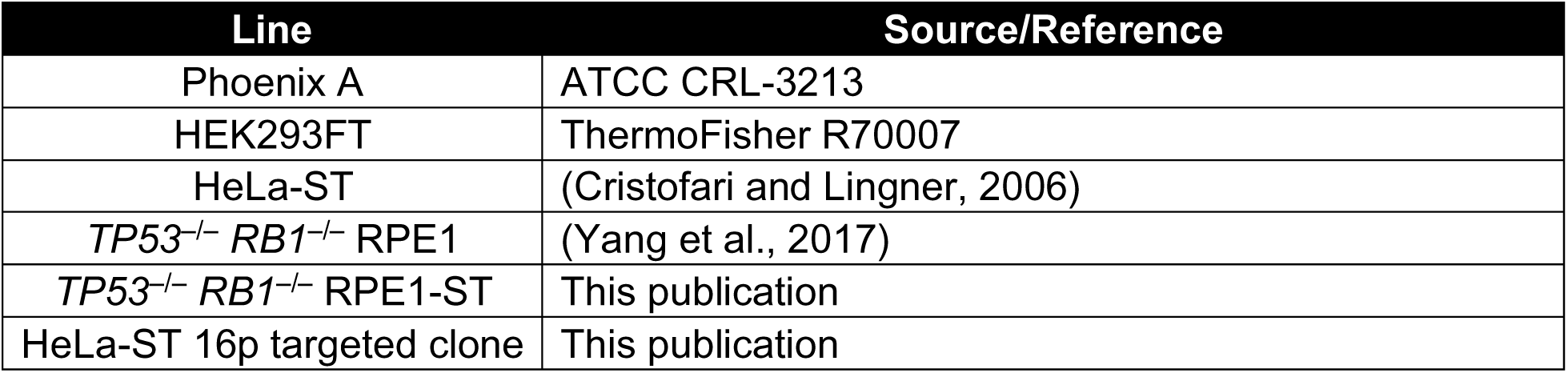

#### shRNA Sequences

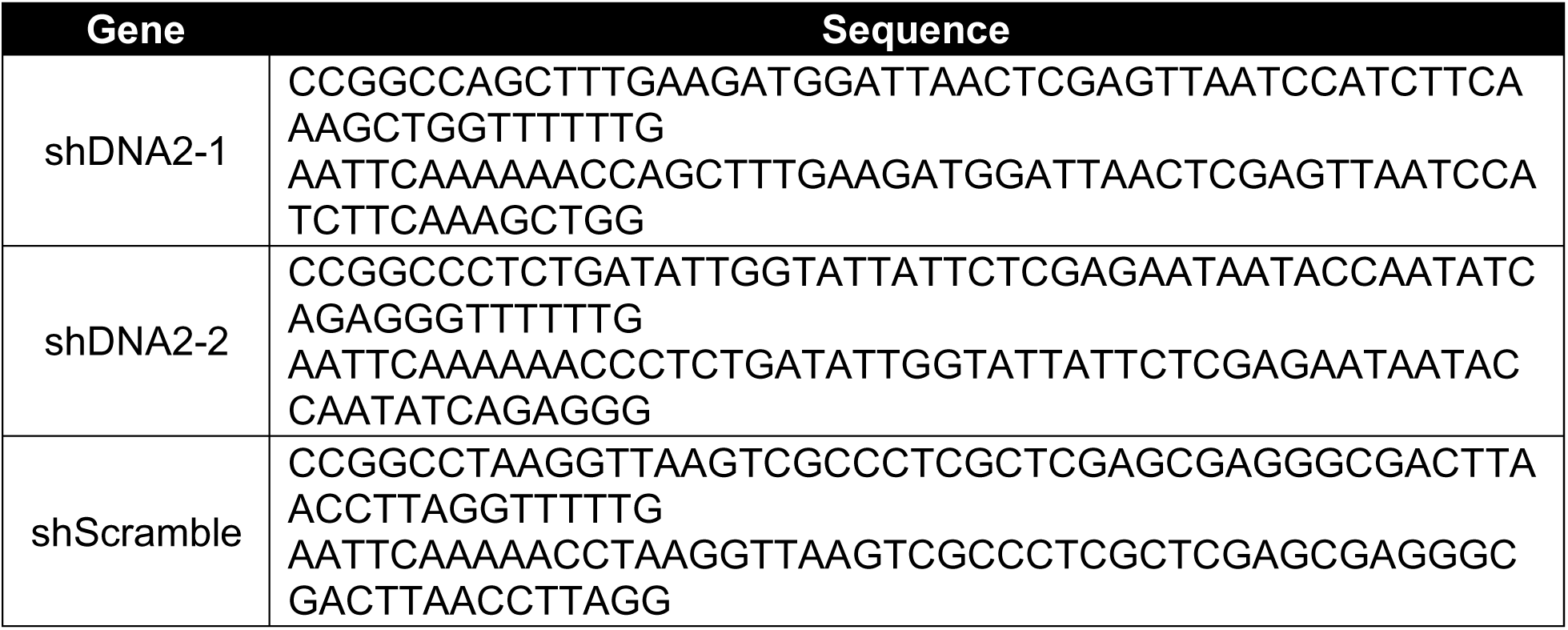

#### siRNAs

| siRNA | Source |
| --- | --- |
| siControl | ThermoFisher 4390843, Silencer Select Negative Control #1 siRNA |
| siRPA70-1 | ThermoFisher 4392420-s12127, Silencer Select RPA70 siRNA |
| siRPA70-2 | ThermoFisher 4392420-s12128, Silencer Select RPA70 siRNA |

#### AsCpf1 crRNA Sequences

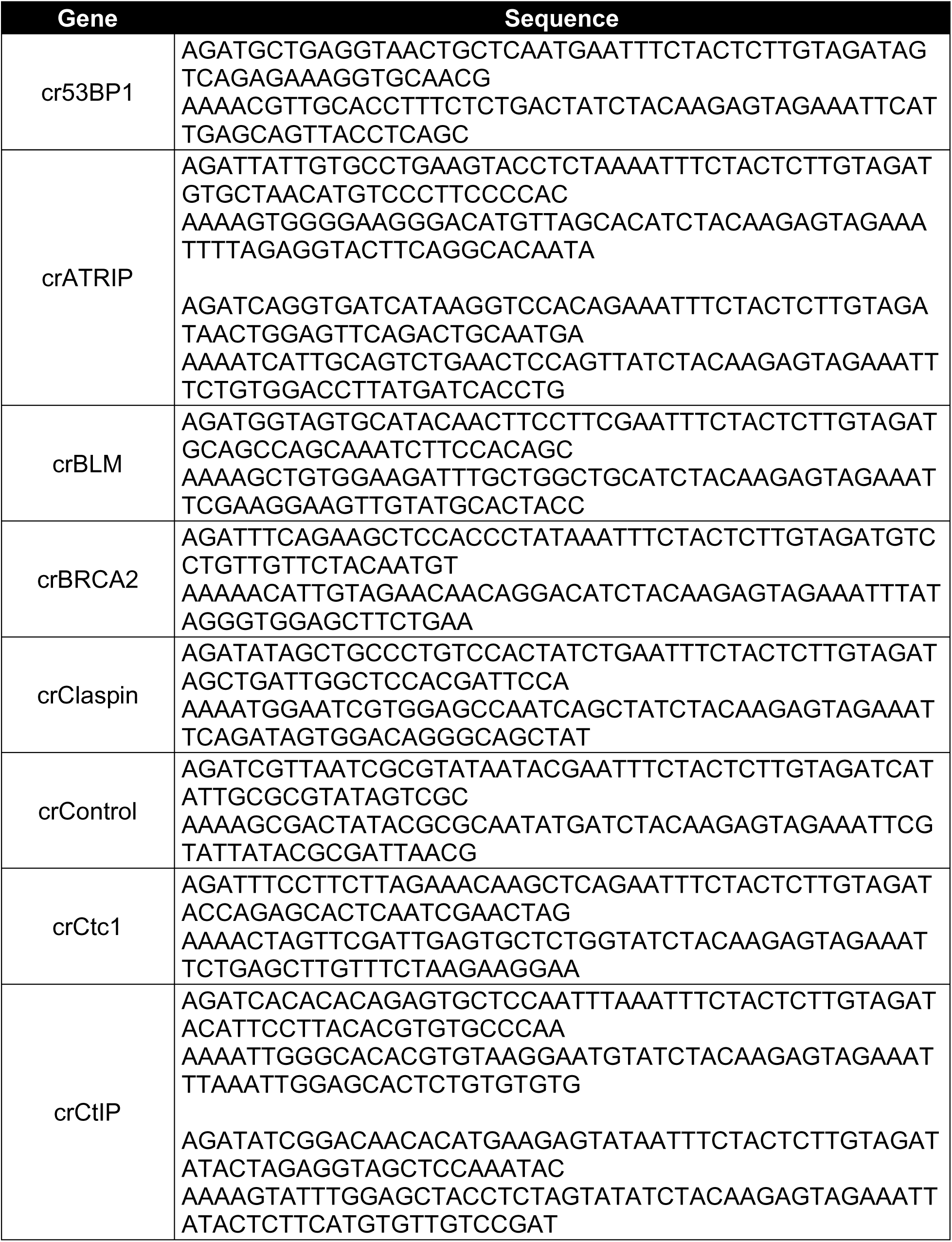
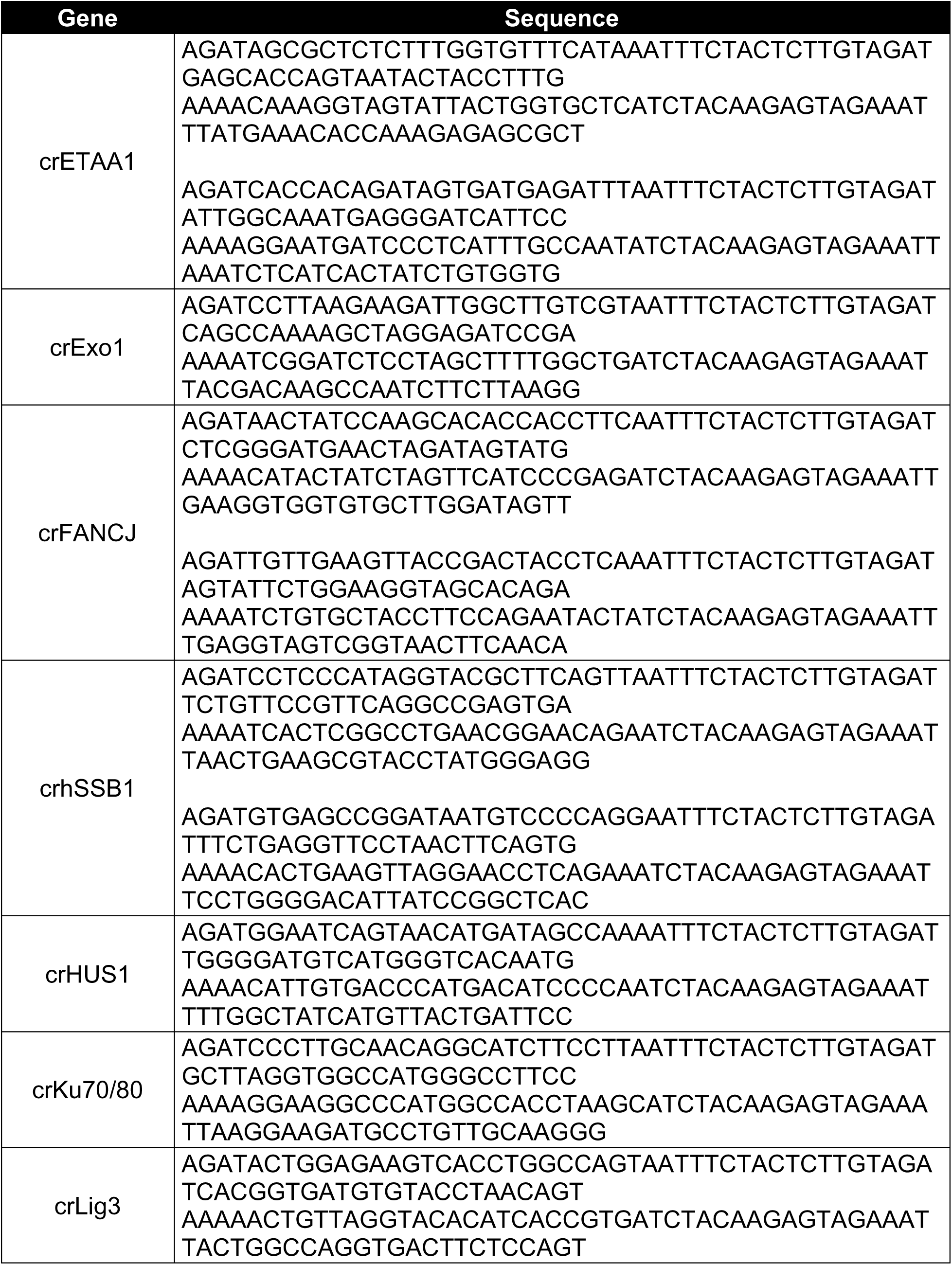
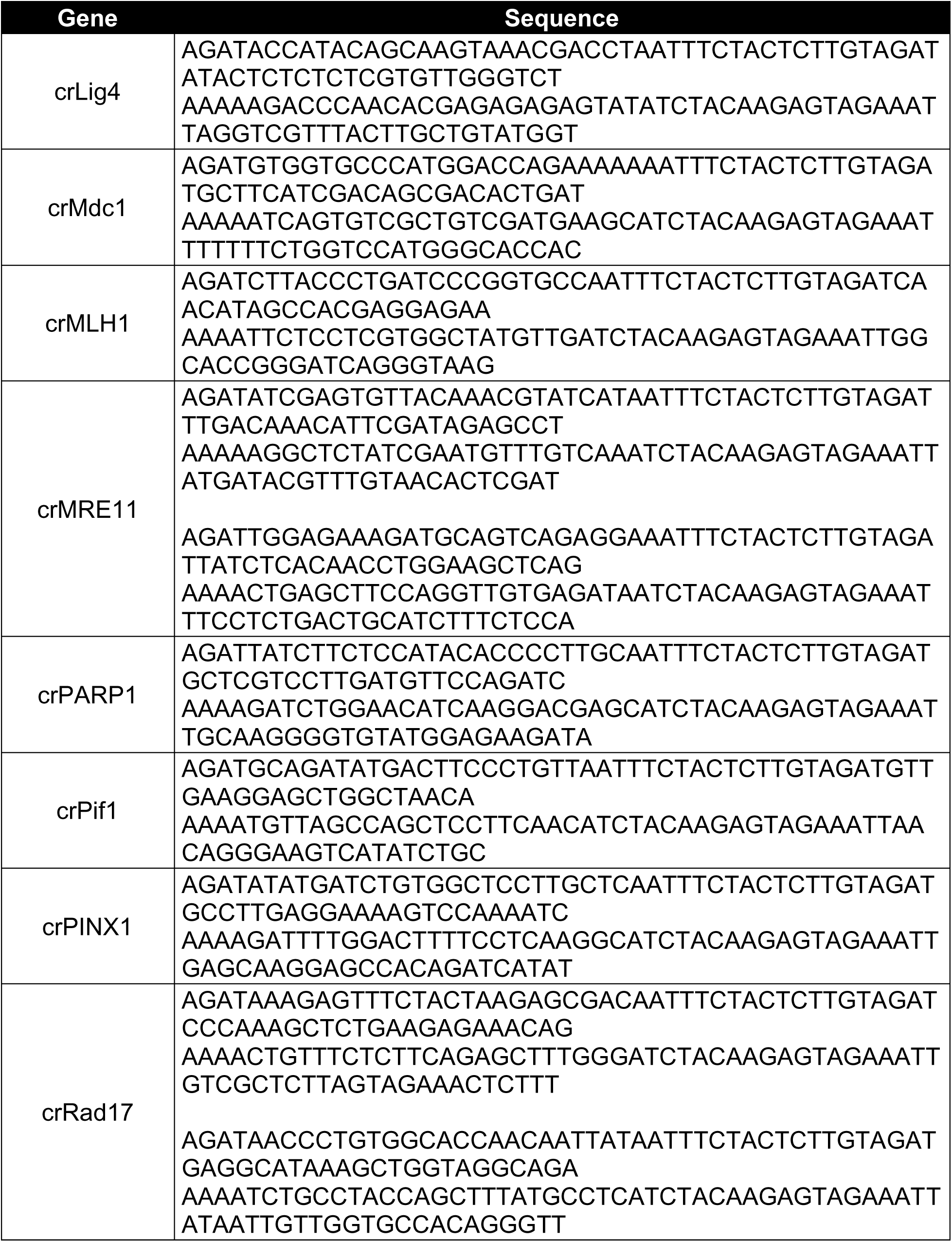
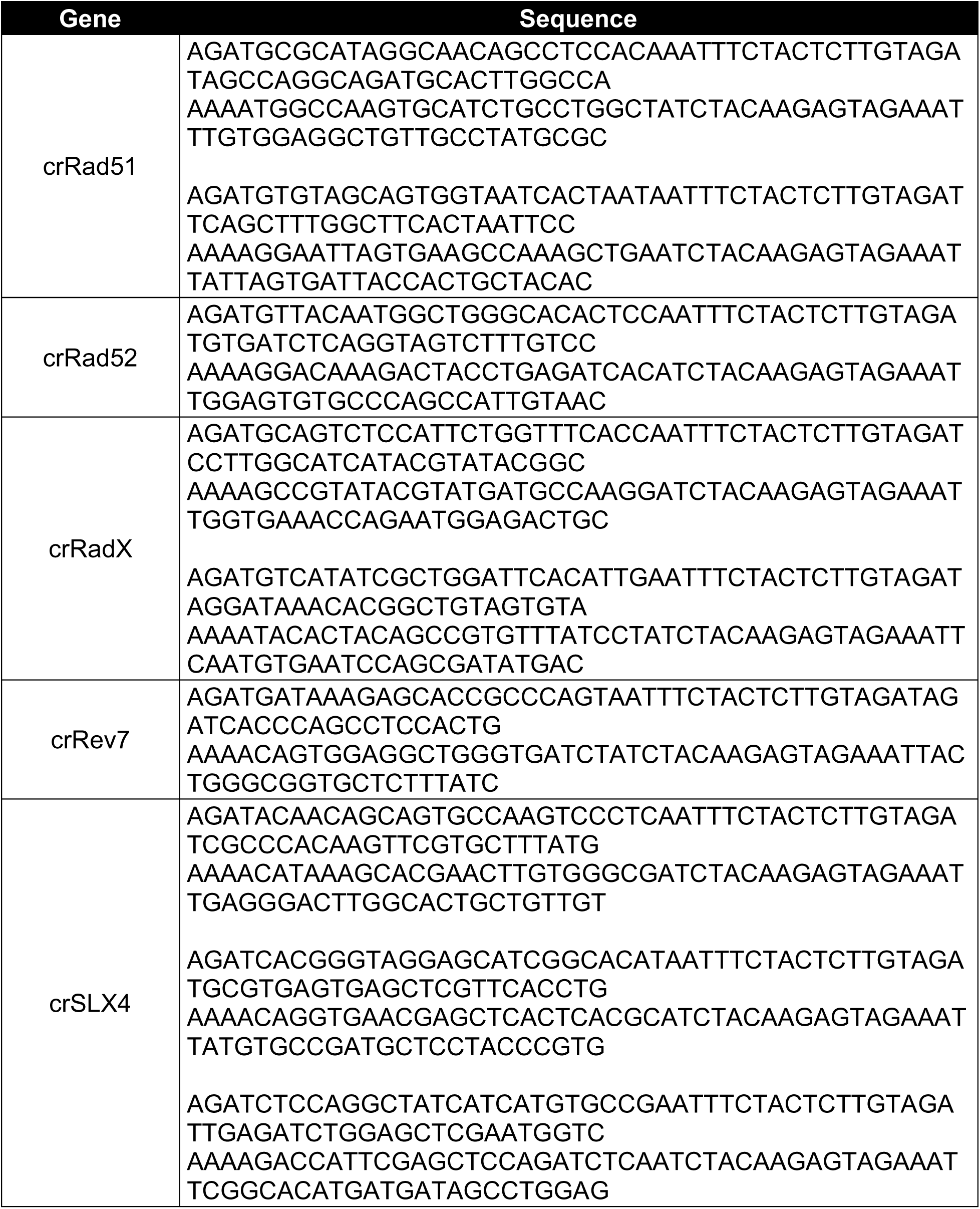
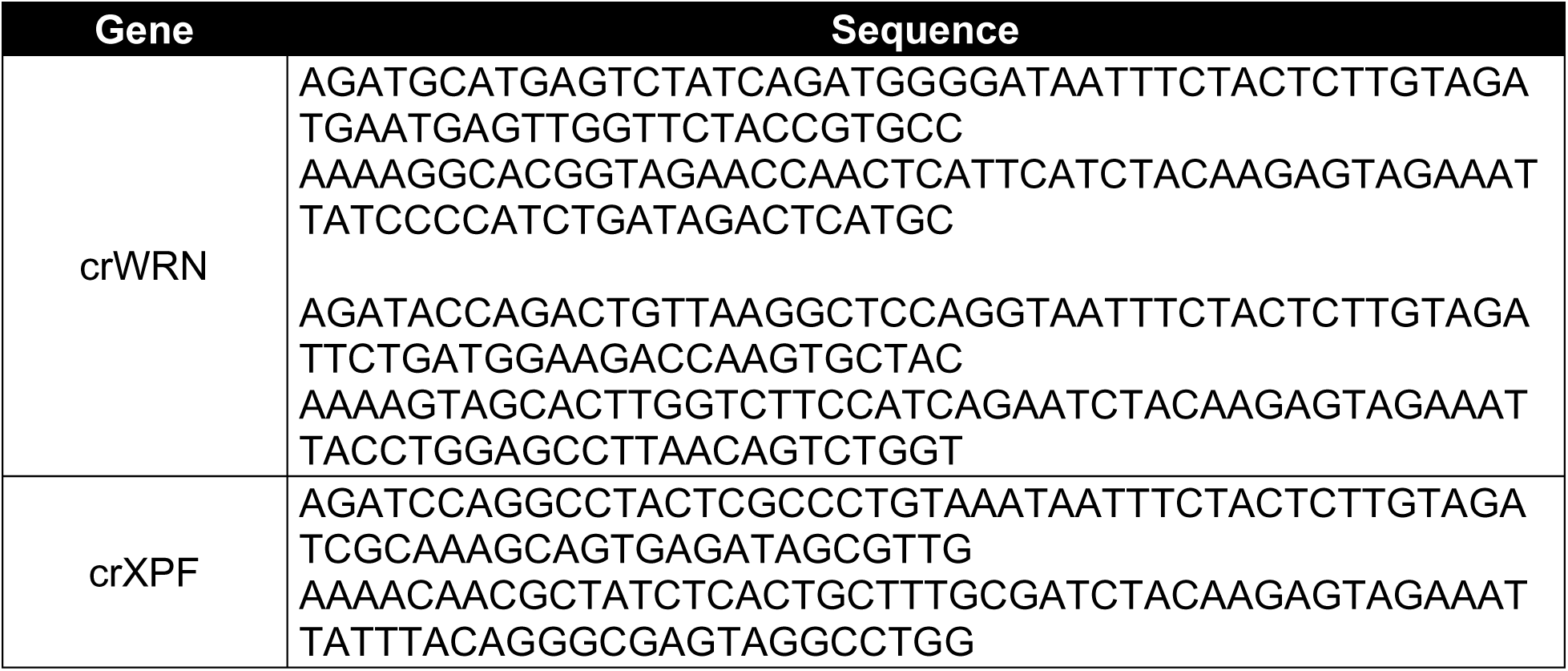

#### Plasmids

| Plasmid | Source |
| --- | --- |
| pBABE-CD hTERT-Puro | (Hahn et al., 1999) |
| pBABE-WT hTERT-U3-hTR-500-Puro | (Wong and Collins, 2006) |
| pET-TaqMan Neotelomere Control | Cloned here |
| pLenti-AsCpf1-Puro | (Zetsche et al., 2017) |
| pLenti-sgRNA | Addgene 71409 |
| pLenti-TaqMan TS-sgLuc-Blast | Cloned here |
| pLenti-TaqMan TS-sgTS-Blast | Cloned here |
| pLKO.1 | Open Biosystems |

#### Antibodies

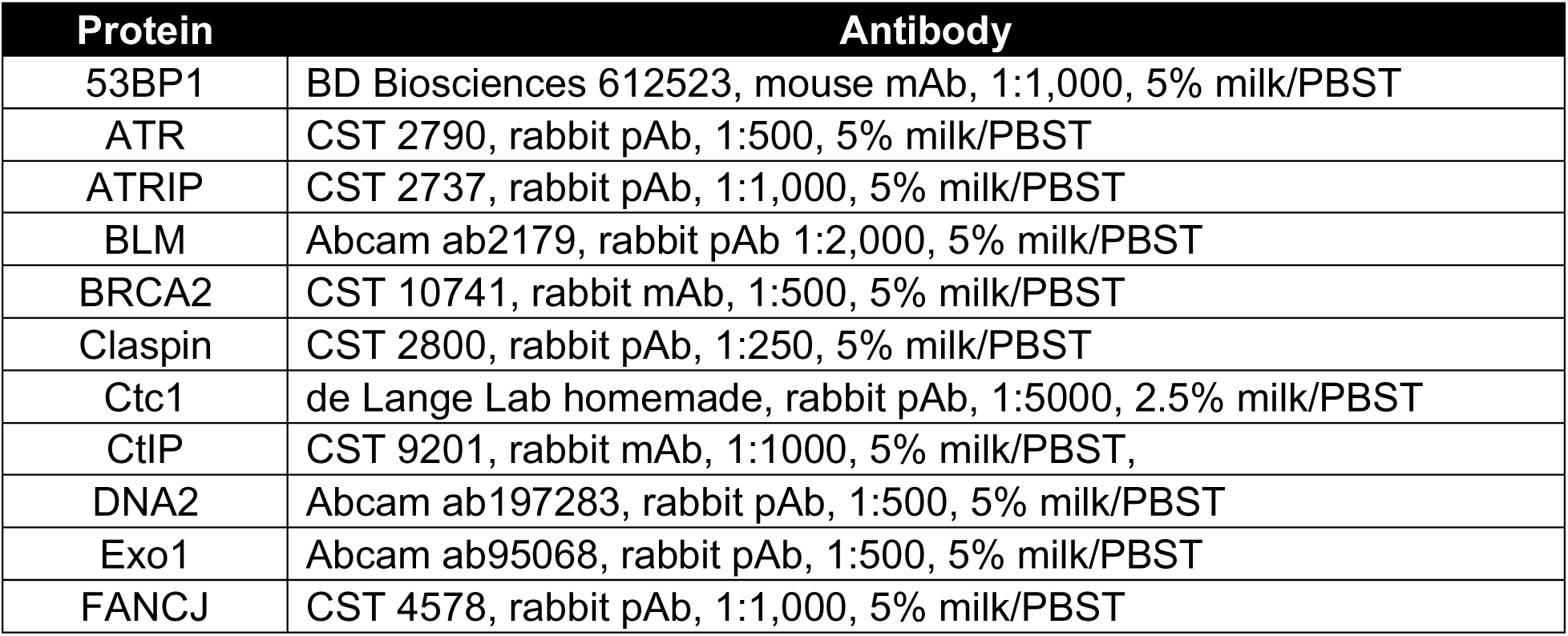
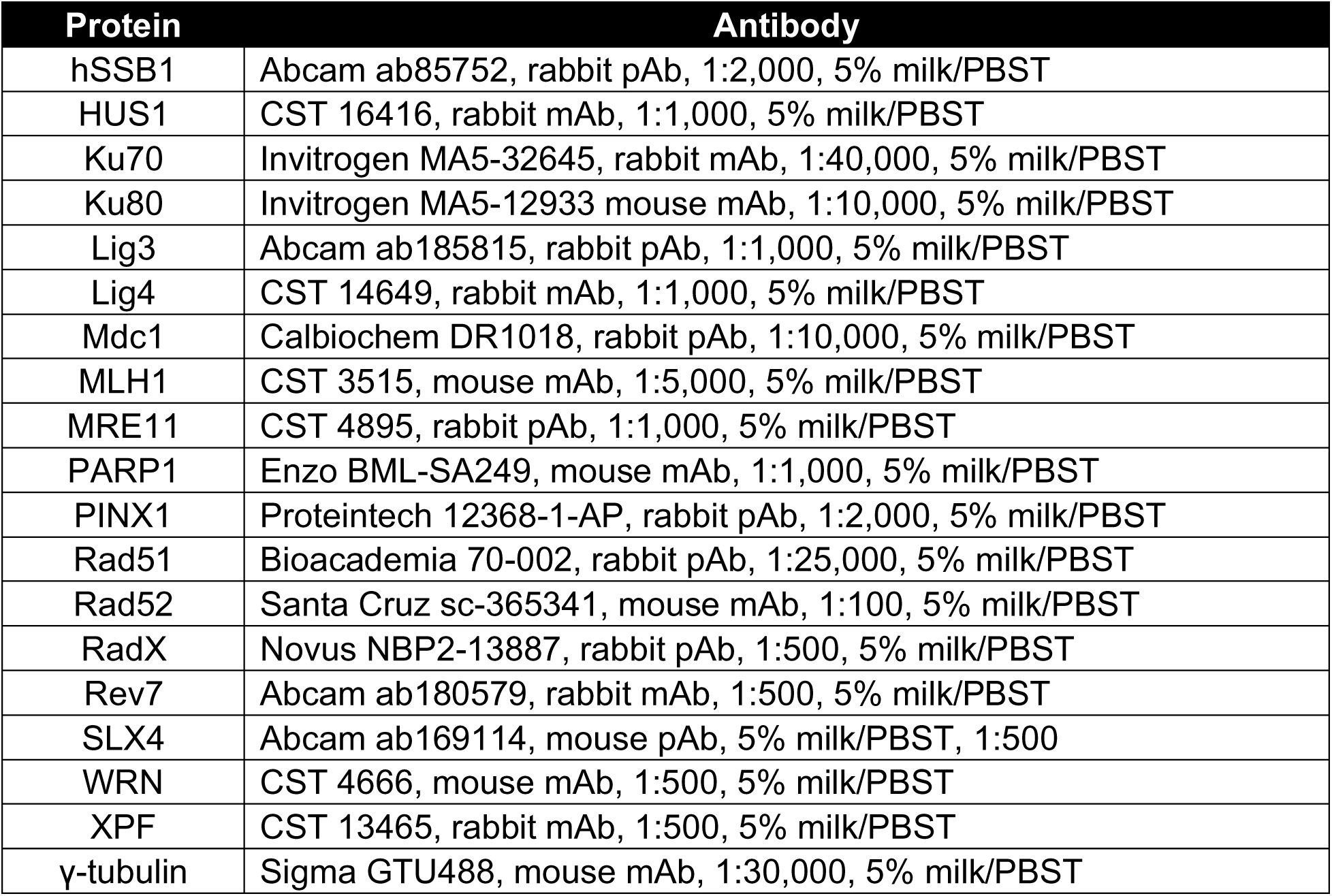

#### Drugs/Chemicals

| Drug/Chemical | Supplier/Solvent |
| --- | --- |
| BIBR1532 | Tocris 2981, DMSO |
| KU55933 | Selleckchem S1092, DMSO |
| AZD-6748 | Selleckchem S8843, DMSO |
| VE-821 | Selleckchem S8007, DMSO |
| M4344 | MedChemExpress HY-136270, DMSO |
| CHIR124 | Selleckchem S2683, DMSO |
| MK-8776 | Selleckchem S2735, DMSO |
| CCT245737 | Selleckchem S8253, DMSO |
| Ganciclovir | Sigma-Aldrich G2536, DMSO |

## Resource availability

### Lead contact

Requests for resources or reagents related to this study may be sent to the Lead Contact, Titia de Lange.

### Materials availability

The unique cell lines, plasmids, and antibodies generated during this study are available upon request.

## Experimental model and subject details

sgTS-TaqMan-TS RPE1 cells were generated by infecting p53^-/-^ Rb^-/-^ RPE1 cells (Yang et al., 2017) with the sgTS-TaqMan-TS lentivirus. RPE1-ST cells were generated by infecting the sgTS-TaqMan-TS RPE1 with a single retrovirus that simultaneously overexpresses hTERT and hTR (Wong and Collins, 2006).

A clonal HeLa-ST cell line (termed HeLa-ST 16p targeted clone, 16p-targ) was derived by integrating the construct shown in Figure 3A into the region between the *LUC7L* and *FAM234A* genes on chromosome 16 (chr16:229,557–229,641). HeLa-ST cells were nucleofected with a linearized template containing diphtheria toxin A cassettes on both ends plus two AsCpf1 plasmids, each encoding a crRNA targeting the region just proximal to the *LUC7L* transcriptional start site: 5′-CCCTCCGTAGAGACT CGTTTGAG-3′ and 5′-GTTCCGCCGTTGGACAACTTGCG-3′. Following nucleofection, cells were selected in 10 µg/mL blasticidin for 10 days, and resistant cells were subcloned by flow sorting. 96 clones were screened in 16-clone pools, and then positive pools were screened individually using the 5′/centromeric and 3′/telomeric PCR primers below. HeLa-ST clone 48 was identified as having an intact cassette at the appropriate locus. This clone was assumed to contain a single insertion.

## Method Details

### Cell culture

HeLa-ST cells were cultured in Dulbecco’s modified Eagle medium (DMEM, Corning) supplemented with 15% fetal bovine serum (FBS, Gibco), non-essential amino acids (Gibco), 2 mM L-glutamine (Gibco), and 100 U/mL penicillin plus 100 μg/mL streptomycin (Gibco). Phoenix A and HEK293FT cells were cultured in DMEM supplemented with 10% bovine calf serum (BCS) and non-essential amino acids, L-glutamine, penicillin, and streptomycin, as above. All RPE1 cell lines were cultured in DMEM/F-12 medium (Gibco) supplemented with 10% FBS (Gibco) and 100 U/mL penicillin plus 100 μg/mL streptomycin (Gibco). All human cells were maintained in a humidified incubator at 37°C and 5% CO_2_.

### Western blotting

Whole-cell lysates were prepared via direct lysis of 10^4^ cells per µL 2X Laemmli lysis buffer (100 mM Tris-Cl pH 6.8, 20% glycerol, 2% SDS, 0.025% bromophenol blue, 300 mM β-mercaptoethanol). Lysed samples were heated at 95°C for 5 minutes and then sonicated. The equivalent of 10^5^ cells was fractionated via SDS-PAGE on Tris-glycine gels and transferred to nitrocellulose membranes. Membranes were blocked in 5% nonfat dry milk in PBST (0.1% Tween-20 in PBS) for 1 hour at room temperature and then incubated with primary antibody using the specified antibody dilution and conditions (see Key Resources Table) overnight at 8°C. Membranes were washed 3 times in PBST, incubated for 1 h at room temperature in secondary antibodies in 5% milk/PBST, and washed 3 times more in PBST. Membranes were developed using chemiluminescence with SuperSignal West Pico PLUS (ThermoFisher).

### Lentiviral and retroviral production and infection

Lentivirus and retrovirus were produced in HEK293FT or Phoenix A cells, respectively as previously described (Wu *et al*., 2012). Two to six infections were performed at 12 h intervals, as required. Cells were selected in an appropriate antibiotic until uninfected control cells were dead.

### sgTS-TaqMan-TS neotelomere formation assay

Human cells in which the frequency of neotelomere formation could be measured were generated via infection with the sgTS-TaqMan-TS lentivirus (or the sgLuc control). The lentiviral plasmid contains the PCR cassette in Figure 1A. This plasmid also directs the constitutive expression of a single-guide RNA (sgRNA) targeting the TS site (or firefly luciferase as a control) under the control of a human U6 promoter. Cas9-mediated cleavage at the TS site was induced by infecting sgTS-TaqMan-TS cells with Cas9 adenovirus (AdCas9; Vector Biolabs). HeLa-ST cells were plated at 0 h at 500,000 cells per 10-cm plate in the presence of 2.0 × 10^7^ PFU AdCas9 adenovirus (MOI = 40) and 4 µg/mL polybrene. RPE1 cells were plated at 0 h at 700,000 cells per 10-cm plate in the presence of 2.8 × 10^7^ PFU AdCas9 (MOI = 40) and 8 µg/mL polybrene. Uninduced control cells were plated at the same density with the same concentration of polybrene but without AdCas9. 24 hours after plating, the media was changed. Genomic DNA was harvested from cells at the indicated intervals. For cells treated with drugs during the assay, drug treatment was initiating on plating with AdCas9 and continued until harvest unless indicated otherwise.

### Genomic DNA extraction

Genomic DNA was isolated from cells using the Zymo Research Quick-DNA Miniprep kit (Zymo Research). Adherent cells were washed once with 1X PBS and then lysed by adding ∼1 mL Genomic Lysis Buffer with 0.5% β-mercaptoethanol. The lysate was scraped into a microcentrifuge tube, vortexed for 30 s, and incubated at room temperature for at least 30 min. Column-based purification of genomic DNA was performed according to the manufacturer’s protocol. Genomic DNA was eluted with sterile ddH_2_O, and its concentration was estimated by UV spectrophotometry (NanoDrop ND-1000). Samples were diluted in sterile ddH_2_O to a final concentration of 25 ng/µL and stored at –20°C.

### Endpoint neotelomere PCR reactions

Endpoint neotelomere PCR reactions were performed with HotStarTaq DNA polymerase (Qiagen). The forward and reverse primer sequences were 5′-AAGTACCCCTATCGCGTGTG-3′ and 5′-ACCCTAACCCTAACCCTAACTCTG-3′, respectively. In general, PCR reactions were performed in a 20-µL volume containing 100 ng genomic DNA template (estimated by UV spectrophotometry, NanoDrop ND-1000), 1X PCR buffer (containing a final MgCl_2_ concentration of 1.5 mM), 500 nM forward primer, 500 nM reverse primer, 200 µM each dNTP, 0.5 U HotStarTaq DNA polymerase. When reaction volumes other than 20 µL were used, the quantities of input genomic DNA and DNA polymerase were scaled accordingly. PCR was carried out on C1000 Touch Themal Cycler (Bio-Rad) device with the following thermocycling conditions: 15 minutes initial denaturation at 95°C, followed by 35 cycles of 94°C for 30 s, 57°C for 30 s, and 72°C for 30 s, with a final extension for 10 minutes at 72°C. Deviations from this themocycling protocol are as specified in the figure legends in the text. PCR products were resolved by agarose gel electrophoresis in 1X TBE and visualized via ethidium bromide staining.

### Neotelomere TaqMan qPCR

The TaqMan qPCR assay used to quantify neotelomere formation was derived from a previously described assay for the detection of cytomegalovirus (CMV) DNA in patient samples (Jebbink et al., 2003). The forward and reverse neotelomere primer sequences were as specified above. The custom neotelomere TaqMan probe had a sequence of 5′-TGGCCCAGGGTACGGATCTTATTCG-3′ and was labeled with 6-carboxyfluorescein (FAM) at the 5′ end as the reporter fluorophore and with Black Hole Quencher 1 (BHQ1) at the 3′ end as the quencher (Biosearch Technologies). qPCR reactions were performed in a 20-µL volume containing 100 ng genomic DNA template (estimated by UV spectrophotometry, NanoDrop ND-1000), 1X AmpliTaq Gold 360 Buffer (ThermoFisher Scientific), 3 mM MgCl_2_, 400 nM forward primer, 400 nM reverse primer, 200 nM TaqMan probe, 200 µM each dNTP, 1 U AmpliTaq Gold 360 DNA polymerase (ThermoFisher Scientific), and 50 nM ROX reference dye (ThermoFisher Scientific). qPCR was carried out on a QuantStudio 12K Flex Real-Time PCR System (Rockefeller University Genomics Resource Center, ThermoFisher Scientific) device with the following thermocycling conditions: 10 minutes initial denaturation at 95°C, followed by 45 cycles of 94°C for 15 s, 60°C for 30 s, and 72°C for 30 s. Fluorescence data were collected during the 72°C step. *C*_T_ values were calculated by the QuantStudio software using a *C*_T_ fluorescence threshold value of 0.192813. At least three technical replicates were performed for each reaction, alongside a ddH_2_O and/or genomic DNA negative control and a 10^−2^ or 10^−3^ neotelomeres per haploid genome positive/standardization control. The absolute number of neotelomeres per haploid genome (*N*) was calculated using the equation *C*_T_ = –3.290log_10_(*N*) + *b*, where the *y*-intercept *b* was calculated for each reaction plate based on the *C*_T_ value calculated for the standardization control. These values were then normalized to the quantity of input genomic DNA calculated based on qPCR with primers targeting genomic *ACTB* (see below).

A standard curve for the neotelomere TaqMan assay was generated with by spiking a TaqMan Neotelomere Control plasmid into 100 ng/µL human genomic DNA at a known copy number. Serial 10-fold dilutions of this mixture were prepared from 10^1^ to 10^−6^ copies of the pET-TaqMan Neotelomere Control plasmid per haploid genome. These standards were used as templates for the TaqMan qPCR reaction described above. The average *C*_T_ values were plotted as a function of the copy number of pET-TaqMan Neotelomere Control per haploid genome (*N*) and fitted to the equation *C*_T_ = *A*log_10_(*N*) + *b* by linear regression in GraphPad Prism 9.0, which yielded *A* = –3.290 and *b* = 25.13 with *R*^2^ = 0.9999.

To normalize to the quantity of input genomic DNA, a parallel qPCR reaction with SYBR Green quantification was performed with primers targeting *ACTB* genomic DNA (NCBI Gene ID 60). The forward and reverse *ACTB* primers were 5′-GTGCTGTGGAAG CTAAGTCCTGC-3′ and 5′-GTCTTTGCGGATGTCCACGTCAC-3′, respectively. qPCR reactions were performed in a 20-µL volume containing 25 ng genomic DNA template (estimated by UV spectrophotometry, NanoDrop ND-1000), 1X SYBR Green PCR Master Mix (ThermoFisher), 500 nM forward primer, and 500 nM reverse primer. qPCR was carried out on a QuantStudio 12K Flex Real-Time PCR System (Rockefeller University Genomics Resource Center, ThermoFisher Scientific) device with the following thermocycling conditions: 10 minutes initial denaturation at 95°C, followed by 35 cycles of 95°C for 15 s and 60°C for 1 minute, followed by melt-curve analysis. Fluorescence data were collected during the 60°C step. *C*_T_ values were calculated by the QuantStudio software using an automatically calculated *C*_T_ fluorescence threshold value. The inferred quantity of input DNA (*Q*) was calculated using the equation *C*_T_ = –3.477log_10_(*Q*) + *b*, where the *y*-intercept *b* was calculated for each reaction plate based on the *C*_T_ value determined for the standardization control.

A standard curve for the *ACTB* qPCR described above was generated using 2-fold serial dilutions of human genomic DNA. The average *C*_T_ values were plotted as a function of the input quantity of DNA (*Q*) and fitted to the equation *C*_T_ = *A*log_10_(*Q*) + *b* by linear regression in GraphPad Prism 9.0, which yielded *A* = –3.477 and *b* = 29.23 with *R*^2^ = 0.9997.

### TRAP assays

Telomeric Repeat Amplification Protocol (TRAP) assays were performed with the TRAPeze Telomerase Detection Kit (Millipore Sigma) essentially according to the manufacturer’s protocol. Briefly, cells were harvested by washing once with 1X PBS, lifting with trypsin-EDTA, and quenching with serum-containing growth medium. 4 × 10^5^ cells were lysed by resuspending in 160 µL 1X CHAPS Lysis Buffer and incubating on ice for 30 minutes. The lysate was cleared by centrifuging at 12,000 *g* for 20 min at 4°C, and the supernatant was transferred to a fresh microcentrifuge tube, giving a lysate containing 2500 cells/µL. Serial fivefold dilutions in 1X CHAPS Lysis Buffer were prepared as necessary. Heat-treated negative controls were prepared by heating the 2500 cells/µL lysate at 95°C for 15 minutes. An additional negative control was performed with 1X CHAPS Lysis Buffer in each trial.

Telomerase-mediated extension and subsequent amplification of TRAP products were conducted in 25-µL reactions containing 1 µL of cell lysate and 2 U of Titanium Taq DNA polymerase (Takara). The other kit components—TRAP reaction buffer, dNTP mix, TS primer, TRAP primer mix, and ddH_2_O—were added in the proportions specified by the manufacturer. Reactions were incubated at 30°C for 30 minutes and 95°C for 2 minutes and then underwent 32 cycles of PCR at 94°C for 15 s, 59°C for 30 s, and 72°C for 1 minute. TRAP assay products were resolved on a 1X TBE 12.5% native polyacrylamide gel at 100 V for 3 hours. Bands were visualized by staining in 1 µg/mL ethidium bromide in ddH_2_O for 30 minutes and then destaining in ddH_2_O for 30 minutes.

### E. coli *exonuclease I digestion*

For samples tested in the *E. coli* exonuclease I (*Exo*I) assays, genomic DNA harvested with the Quick-DNA Miniprep kit (Zymo Research) was digested with 100 µg/mL RNase A in 1X TNE for 3 hours and then with 100 µg/mL proteinase K in 1X TNES. Genomic DNA was extracted with phenol-chloroform-isoamyl alcohol and precipitated with isopropyl alcohol overnight. 15 µg genomic DNA was digested with 60 U *E. coli Exo*I (New England BioLabs M0293) in 1X *Exo*I buffer (67 mM glycine-NaOH pH 9.5, 6.7 mM MgCl_2_, 10 mM β-mercaptoethanol) overnight at 37°C. After the overnight digest, the reactions were chased with another 60 U *Exo*I for 2 additional hours at 37°C. As a negative control, 15 µg genomic DNA was incubated overnight in 1X *Exo*I buffer without enzyme. Genomic DNA was extracted, precipitated, and used as input for the Neotelomere TaqMan qPCR assay and overhang gels.

### In-gel analysis of single-stranded telomeric DNA

After *Exo*I treatment, genomic DNA samples were digested overnight with MboI (New England BioLabs) at 37°C according to the manufacturer’s specifications. Following digestion, DNA concentrations were measured by Hoechst fluorimetry, and 1.5 µg DNA was resolved by agarose gel electrophoresis in 0.5X TAE. In-gel hybridization with a γ-^32^P-ATP end-labelled (AACCCT)_4_ probe was performed as previously described (Mirman *et al*., 2022) under native conditions to detect single-stranded telomeric overhangs and then after in situ denaturation to detect total telomeric DNA.

### AsCpf1 knockouts

Bulk knockout by AsCpf1 was achieved by infecting cells four times with pLenti-AsCpf1 lentiviruses (Zetsche *et al*., 2017) containing the oligonucleotide crRNA cassettes listed in the Key Resources Table. In scenarios where a satisfactory bulk knockout could not be achieved with a single crRNA cassette, cells were co-infected with pLenti-AsCpf1 lentiviruses encoding distinct crRNAs.

### Pif1 RT-qPCR

Total RNA was isolated from a 10-cm plate of RPE1 cells using TRIzol (ThermoFisher) according to the manufacturer’s protocol. RNA was digested with RNase-free DNase I (Roche) to remove contaminating genomic DNA according to the manufacturer’s protocol, reextracted with TRIzol, and precipitated once again with isopropyl alcohol overnight. cDNA was synthesized with SuperScript IV reverse transcriptase (ThermoFisher) using 1 µg of total RNA and the (dT)_20_ primer. The RNA-primer mixture was pre-annealed by heating to 65°C for 5 minutes and then snap-cooling on ice for at least 1 minute. Reverse transcription was carried out with the following thermocycling program: 10 minutes at 50°C, 5 minutes at 55°C, and 10 minutes at 80°C. cDNA was diluted 25-fold for qPCR.

qPCR reactions were performed with SYBR Green PCR Master Mix (ThermoFisher) to detect Pif1 cDNA and GAPDH cDNA as a normalization control. The forward and reverse Pif1 primers were 5′-AGGAGCTGCCAGGTAAGGTA-3′ and 5′-GCTAACAGGACACTGGGCAT-3′, respectively. The forward and reverse GAPDH primers were 5′-AGCCACATCGCTCAGACAC-3′ and 5′-GCCCAATACGACCAAATCC-3′, respectively. qPCR reactions were performed in a 20-µL volume containing 1 µL 1-to-25 diluted cDNA, 1X SYBR Green PCR Master Mix (ThermoFisher), 500 nM forward primer, and 500 nM reverse primer. qPCR was carried out on a QuantStudio 12K Flex Real-Time PCR System (Rockefeller University Genomics Resource Center, ThermoFisher Scientific) device with the following thermocycling conditions: 10 minutes initial denaturation at 95°C, followed by 40 cycles of 95°C for 15 s and 61°C for 1 minute, followed by melt-curve analysis. Fluorescence data were collected during the 61°C step. *C*_T_ values were calculated by the QuantStudio software using an automatically calculated *C*_T_ fluorescence threshold value. Pif1 qPCR reactions that had more than one melting-curve peak were discarded. The amplification factor for the Pif1 qPCR reaction was determined to be 1.8845, and that for the GAPDH qPCR reaction was assumed to be 2. The quantity of Pif1 mRNA was normalized to the quantity of GAPDH mRNA.

### shRNA knockdown

shRNA-mediated knockdown was achieved by infecting cells with pLKO.1 lentiviruses (Open Biosystems) containing the oligonucleotide cassettes listed in the Key Resources Table. In scenarios where a satisfactory knockdown could not be achieved with a single shRNA, cells were co-infected with pLKO.1 lentiviruses encoding distinct shRNAs.

### siRNA knockdown of RPA70

48 hours prior to Infection with AdCas9, 500,000 RPE1-ST-1 cells previously infected with the TaqMan-TS-sgTS lentivirus were reverse-transfected with Silencer Select siRNAs (ThermoFisher) in using Lipofectamine RNAiMAX (ThermoFisher). 48 hours later, one of three replicate transfected plates was trypsinized, counted, and harvested for Western blotting. On one of the two remaining plates, the media was changed to RPE1 media plus 8 µg/mL polybrene. On the other remaining plate, Ad-Cas9 adenovirus was added to an MOI of 40 in RPE1 media containing 8 µg/mL polybrene. Plates were harvested for genomic DNA 96 hours after transfection.

### Isolation of HeLa-ST clones containing stable neotelomeres

The HeLa-ST 16p targeted clone was infected with pLenti-sgTS-Puro and selected with 0.6 µg/mL puromycin to instate constitutive expression of an sgRNA targeting the integrated TS site. 500,000 cells were infected with 2.83 × 10^7^ PFU Ad-Cas9 adenovirus (MOI = 56.7) in 8 µg/mL polybrene. 24 hours later, the media was changed, and the cells were treated with another 5.67 × 10^7^ PFU Ad-Cas9 adenovirus in media containing 16 µg/mL polybrene. 10 days after the first Cas9 adenovirus infection, the cells were split into 15-cm plates at 5,000 cells per plate in 50 µM ganciclovir (Sigma). Surviving colonies were isolated with cloning cylinders and expanded in 50 µM ganciclovir. Cells were screened for potential neotelomere formation via PCR with primer pairs to detect telomeric repeat addition at the TS site, retention of the 5′/centromeric junction of the TS cassette, and loss of the 3′/telomeric junction of the TS cassette. The primers used to detect telomeric repeat addition were the same as those used in the neotelomere TaqMan qPCR assay. The forward and reverse primers were 5′-AAGTATATGCCAGACTATGCACACA-3′ and 5′-CGCAGGCGCATAACATCAAA-3′, respectively, for the 5′/centromeric junction and 5′-CGGCTCCATACCGACGATAT-3′ and 5′-GACTTCTCTCAGGCAGGCG-3′, respectively, for the 3′/telomeric junction. PCR was carried out with Q5 High-Fidelity 2X Master Mix (New England BioLabs) using the conditions recommended by the manufacturer for 35 cycles, except as specified below. In all reactions, each primer was present at a concentration of 500 nM, and extension was conducted at 72°C for 35 s. Annealing was conducted for 30 s at 68°C for the neotelomere primers, 66°C for the 5′/centromeric junction primers, and 67°C for the 3′/telomeric junction primers. PCR products were resolved on 1X TBE agarose gels with ethidium bromide staining.

### Metaphase FISH

BAC probes were purchased from BACPAC Genomics to identify the 16q arm (Chr16q13, RP11-109J21) and distal p arm of chromosome 16 (Chr16p13.3, RP11-344L06). Preparation of metaphases and probe labeling by nick-translation were performed as previously described (Dewhurst *et al*., 2021), except probes were denatured at 76°C for 10 minutes rather than at 80°C for 8 minutes. Slides were treated with 50 µg/mL RNase A in 2X SSC at 37°C for 10 minutes and then rinsed with 1X PBS. Thereafter, slides were crosslinked in 4% formaldehyde in 1X PBS for 5 min, rinsed thrice with 1X PBS, and dehydrated in an ethanol series. After preheating to 65°C for 1 hour, slides were denatured at 76°C for 110 s in 70% formamide in 2X SSC and then dehydrated in a chilled ethanol series. Slides were hybridized with labeled BAC probes overnight in a humidified chamber at 37°C and washed thrice with 1X SSC at 65°C for 5 minutes each time and then twice with 0.1X SSC at 50°C for 5 minutes each time. All subsequent steps were performed as previously described (Dewhurst *et al*., 2021), except the following antibodies were used at a 1:1000 dilution to detect the FISH probes: AlexaFluor 488 mouse monoclonal anti-digoxin (JacksonImmunoResearch 200-542-156) and AlexaFluor 594 mouse monoclonal anti-biotin (JacksonImmunoResearch 200-582-211).

### Southern blotting for the TS cassette

Genomic DNA was harvested from cells lysed in TNES containing 50 µg/mL proteinase K overnight at 37°C, then extracted with phenol-chloroform-isoamyl alcohol, and precipitated with isopropyl alcohol. Genomic DNA was subsequently digested with 100 µg/mL RNase A in 1X TNE for 4 hours at 37°C, and then an equal volume of TNES containing 100 µg/mL proteinase K was added. DNA was re-extracted with phenol-chloroform-isoamyl alcohol and precipitated with isopropyl alcohol. Samples were digested overnight with PvuII-HF (New England BioLabs 3151) at 37°C according to the manufacturer’s specifications. DNA concentrations were measured by Hoechst fluorimetry, and 8 µg DNA was resolved by agarose gel electrophoresis in 0.5X TAE. The gel was depurinated in 0.25 M HCl for 30 minutes, denatured with 1.5 M NaCl and 0.5 M NaOH twice for 30 minutes each time, and then neutralized with 3 M NaCl and 0.5 M Tris-Cl pH 7.0 twice for 30 minutes each time. Samples were blotted onto a Hybond membrane (GE Healthcare) in 20X SSC. The membrane was UV-crosslinked in a Stratalinker and prehybridized for 1 hour at 65°C with Church mix (0.5 M sodium phosphate buffer at pH 7.2, 1 mM EDTA, 0.7% SDS, 0.1% BSA). A Klenow α-^32^P-dCTP-labeled probe was prepared from a gel-purified 904-bp PCR product spanning the ψ, *gag*, and part of the TS sequence, as indicated in Figure 3A. Before hybridization with the membrane, 2 µg MboI-digested human genomic DNA was added to the probe, and the mixture was heated to 95°C for 5 minutes and then incubated at room temperature for 10 minutes to reduce background. The membrane was hybridized with this mixture in Church mix overnight at 65°C, washed thrice for 15 min each time with Church wash (40 mM sodium phosphate buffer at pH 7.2, 1 mM EDTA, 1% SDS) and then exposed to a PhosphorImager screen overnight.

### T7 endonuclease I assay

PCR products were amplified from the specified genomic DNA samples with Q5 High-Fidelity 2X Master Mix (New England BioLabs) according to the manufacturer’s protocol. The forward and reverse primers were 5′-GACGTTGGGTTACCTTCTGC-3′ and 5′-TCCGATCGCGACGATACAAG-3′, respectively, and the annealing temperature was 67°C for 30 s. PCR products were purified with the QIAquick PCR Purification Kit (Qiagen) and eluted in 10 mM Tris-HCl pH 8.0. 200 ng of the purified PCR product was digested with T7 endonuclease I (New England BioLabs) as specified by the manufacturer. Reactions were quenched with EDTA and then resolved on 1X TBE agarose gels with ethidium bromide staining.

## Quantification and Statistical Analysis

The number of biological replicates performed is indicated in each Figure and corresponds to the number of data points plotted. All metaphase FISH images were blinded before scoring. Statistical analysis was performed in GraphPad Prism. Data plotted throughout are mean ± standard deviation (SD). The level of statistical significance was indicated by asterisks: ns *p* > 0.05, * *p* < 0.05, ** *p* < 0.01, *** *p* < 0.001, **** *p* < 0.0001. Two-way comparisons were performed via two-tailed unpaired or two-tailed ratio-paired *t*-tests, as indicated in the figure legends. As a rule, two-tailed ratio-paired *t*-tests were performed to determine whether the fold-change resulting from a given intervention was significantly different from 1.

## Supplemental Figures and Legends

**Figure S1.**
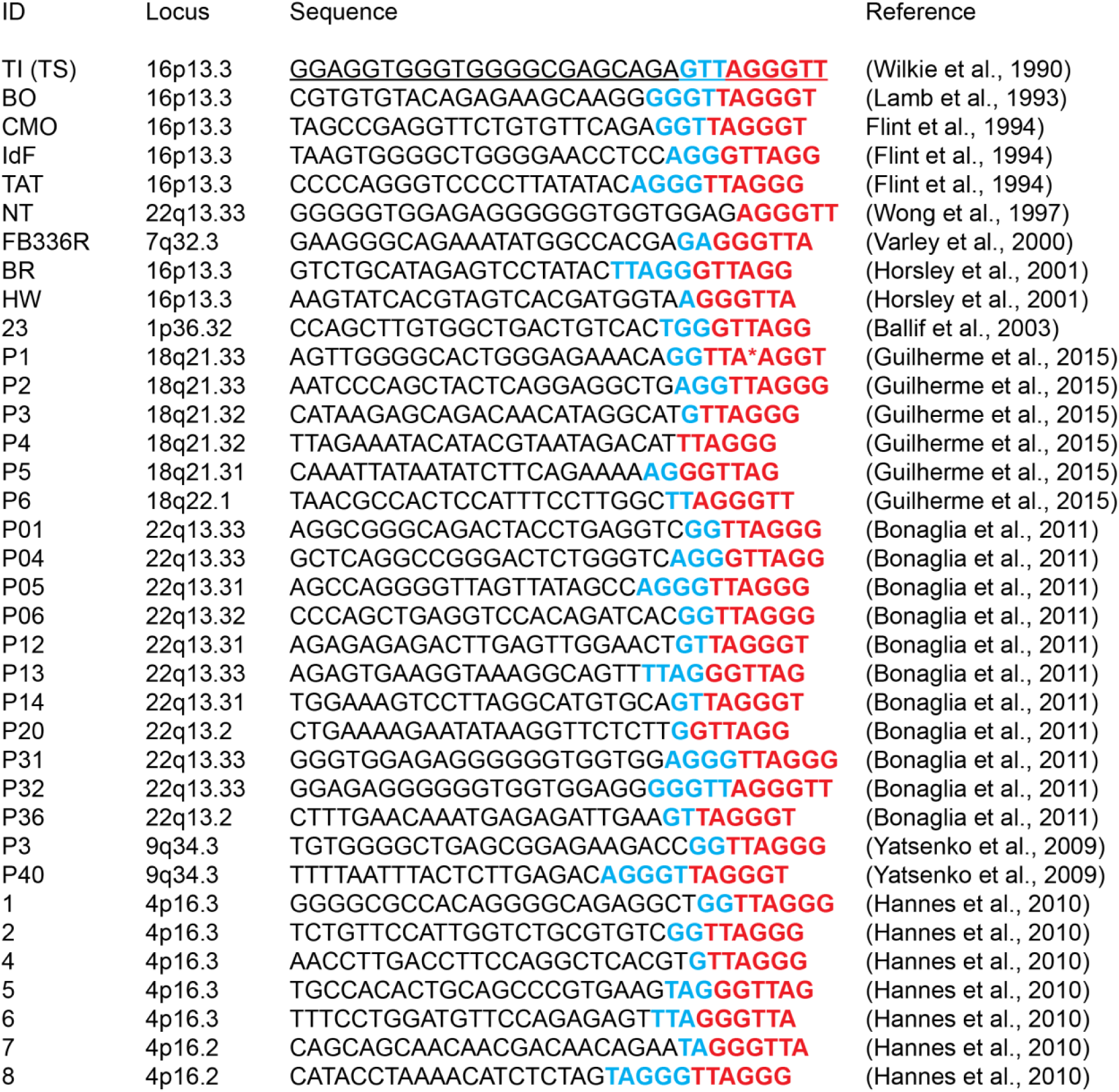
Breakpoint sequences from suspected neotelomere formation events in patients with terminal chromosomal deletions, related to Figure 1 Candidate neotelomere formation events were culled from the literature by identifying terminal deletions in which telomeric repeats are juxtaposed with reference genomic sequence and microhomology to telomeric repeats can be identified at the breakpoint. The 25-nt sequence proximal to unambiguous telomeric sequence—that is, telomeric sequence not present in the reference genome—is provided followed by the first telomeric repeat in red, bold text. Microhomology to telomeric sequence is indicated by blue, bold text. No microhomology is present at the breakpoint in patients NT and P4. TS is derived from patient TI (underlined, top). *In patient P1, telomerase appears to have misincorporated an A instead of a G. Distal telomeric repeats in P1 have the correct sequence.

**Figure S2.**
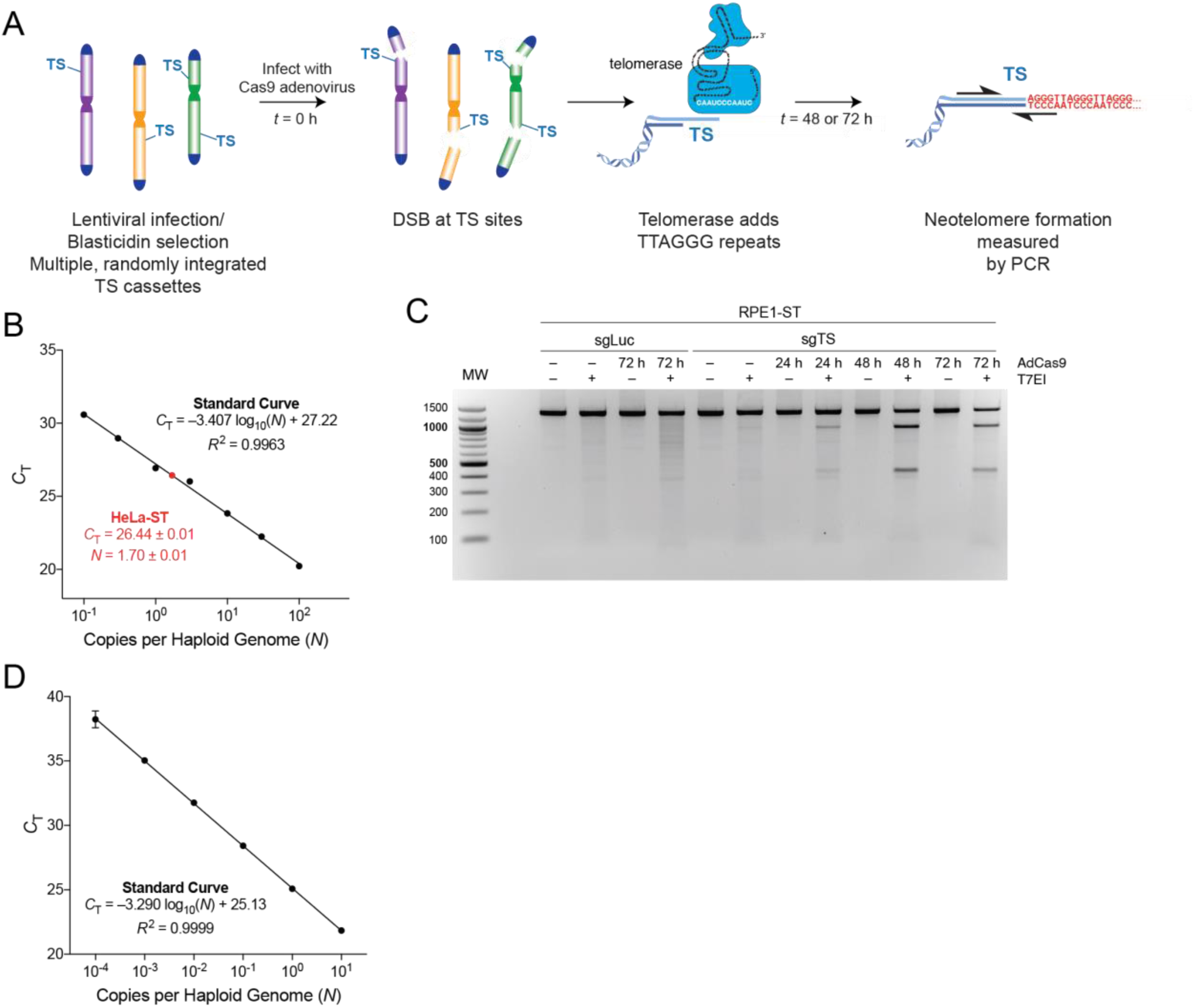
The TaqMan neotelomere formation assay workflow and operating characteristics, related to Figure 1 (A) Schematic depicting the strategy for detecting neotelomere formation. (B) Standard curve for copy number measurement of the sgTS-TaqMan-TS cassette using a plasmid template spiked into human genomic DNA at a known copy number. The data plotted in black are the mean *C*_T_ of three technical replicates versus the base-10 logarithm of the number of plasmid template copies per haploid genome. The data were fitted by linear regression in GraphPad Prism 9 to obtain the displayed equation. The red data point corresponds to the mean *C*_T_ value and the calculated number of integrated sgTS-TaqMan-TS cassettes in HeLa-ST cells per haploid genome from three biological replicates. (C) Ethidium-bromide-stained agarose gel showing the result of a T7 endonuclease I assay performed on PCR products corresponding to the TS site in RPE1 cells expressing sgLuc or sgTS at the indicated times after infection with Cas9 adenovirus. T7 cleavage products are expected at ∼400 and ∼1000 bp. (D) Standard curve for the TaqMan neotelomere formation assay using a positive control plasmid template spiked into human genomic DNA at a known copy number. Three independent qPCRs were performed, each in technical quadruplicate. The data plotted are mean *C*_T_ ± SD versus the base-10 logarithm of the number of template copies per haploid genome. Most error bars are too small to be displayed. The data were fitted by linear regression in GraphPad Prism 9 to obtain the displayed equation.

**Figure S3.**
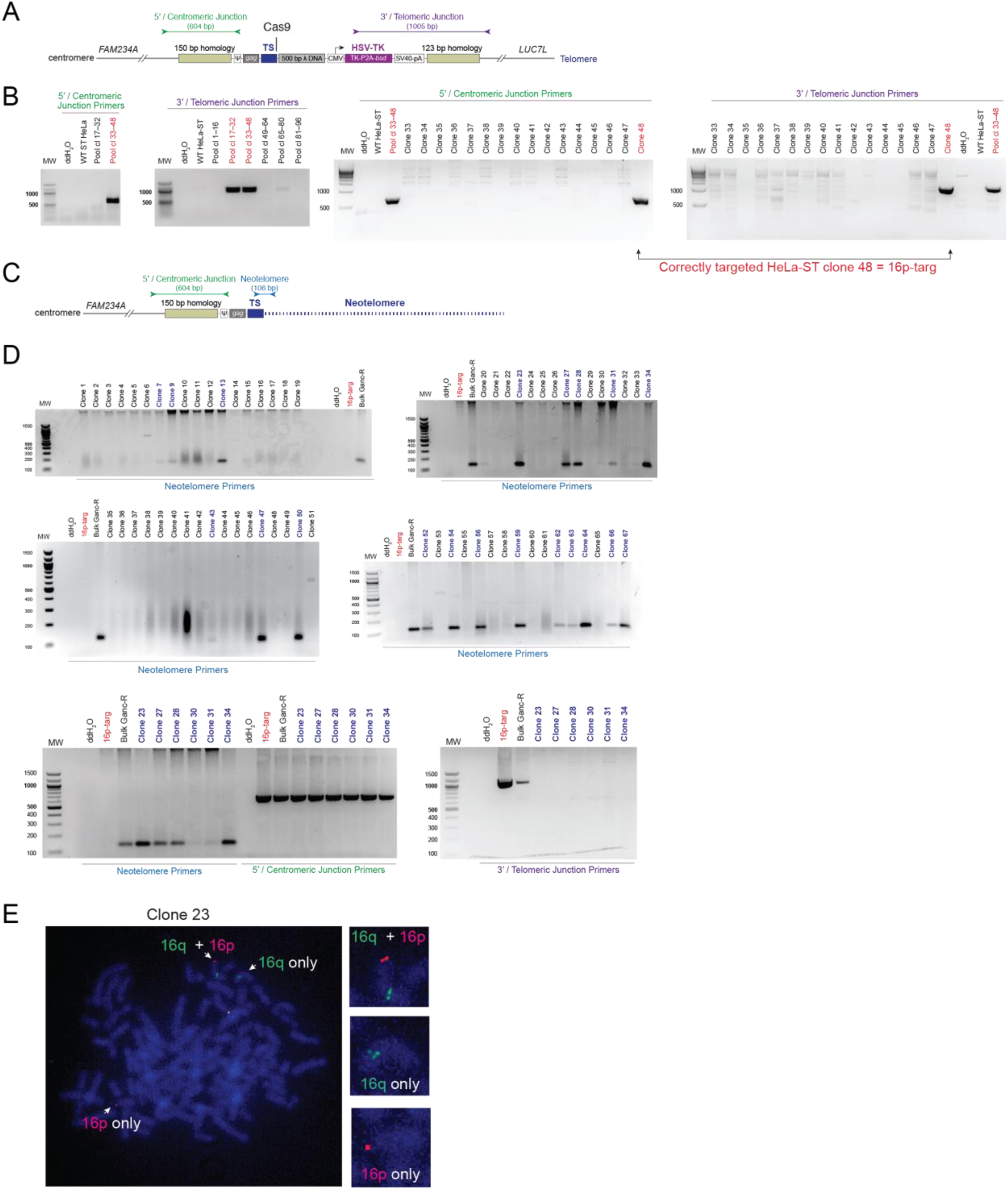
Identification of candidate HeLa-ST clones bearing putative neotelomeres at a predetermined locus, related to Figure 3 (A) Schematic depicting the CRISPR/Cas12a-edited chromosome 16 in the HeLa-ST 16p targeted clone (16p-targ) and the PCR-based strategy to identify a HeLa-ST clone containing the knock-in cassette at the appropriate location. PCR primers were designed to target amplicons spanning the 5′/centromeric and 3′/telomeric junctions between the endogenous chromosome 16 and knock-in cassette. Abbreviations are as in Figure 3A. (B) Ethidium-bromide-stained agarose gels showing PCR products obtained from targeted HeLa-ST pools and clones to identify a clone that contained the TS insert at the 16p locus. Single-cell clones were screened first in pools and then individually with the two primer pairs in (A) to identify clones yielding the expected centromeric and telomeric junction PCR products. Clone 48 (red) was the only clone identified as containing the correct insertion at both ends of the knock-in cassette. This clone was renamed HeLa-ST/16p-targ. (C) Schematic depicting the predicted map of the edited copy of chromosome 16 in HeLa-ST/16p-targ following successful neotelomere formation at the TS site. The binding sites of primer pairs designed to screen for 5′/centromeric junction retention and neotelomere formation are denoted. (D) Ethidium-bromide-stained agarose gels from PCR assays used to identify HeLa-ST/16p-targ derived clones with neotelomeres at TS. Genomic DNA was harvested from the indicated Ganc-R clones and PCR-amplified with the indicated primer pairs shown in (A) and (C). Sterile water and genomic DNA from the parental HeLa-ST/16p-targ (16p-targ) were used as negative controls. Genomic DNA derived from a mixed population of ganciclovir-resistant (Ganc-R) clones was used as a positive control. A subset of clones was further analyzed for retention and loss of the expected segments at the targeted locus (bottom gels). Neotelomere clones are highlighted in bold, blue type. (E) Representative metaphase from Ganc-R clone 23 hybridized with FISH probes for chromosome 16q and 16p (see Figure 3) showing the presence of one intact chromosome 16, one chromosome with the 16q signal (consistent with neotelomere formation), and one chromosome with the 16p signal.

**Figure S4:**
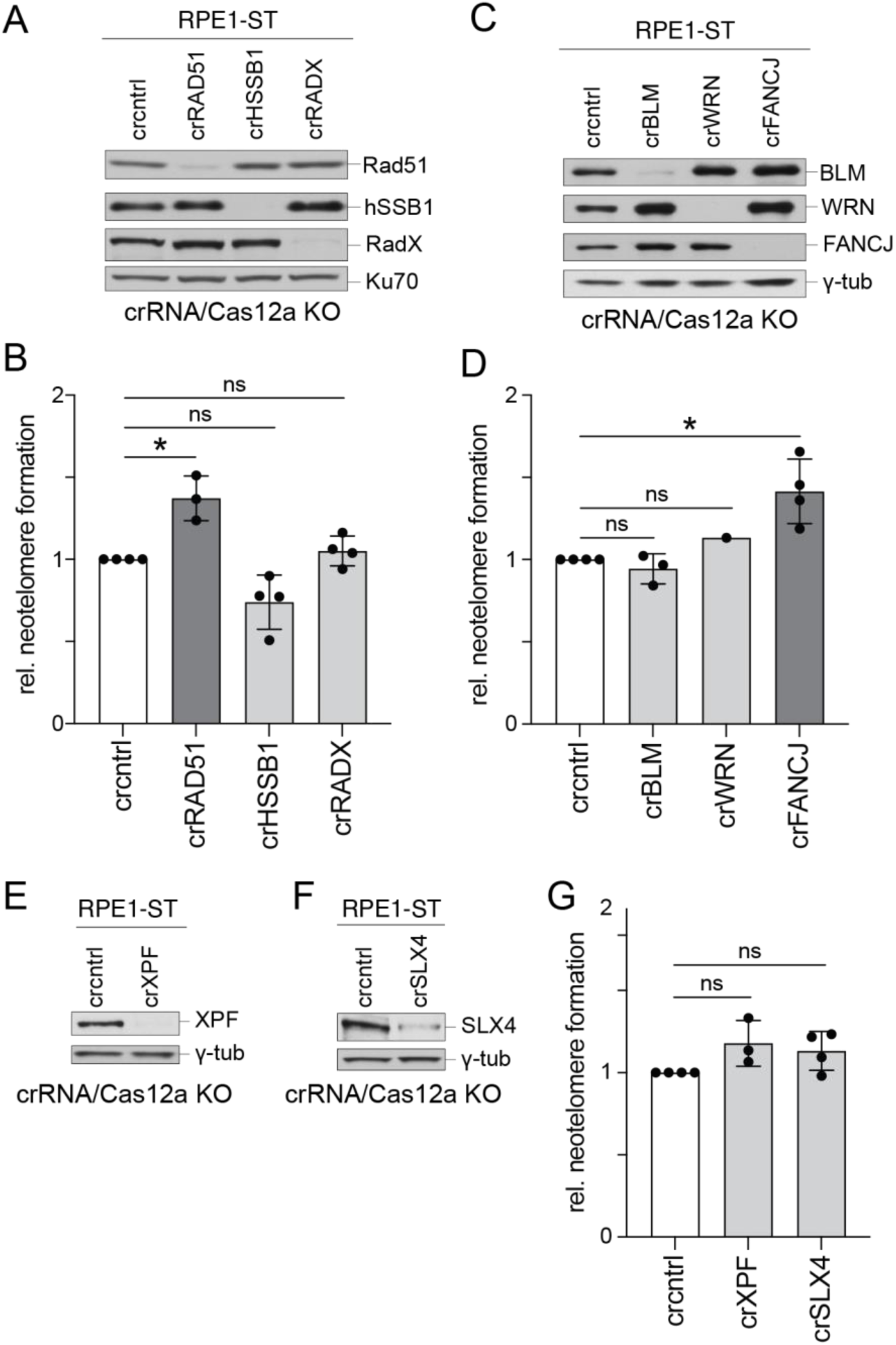
Targeted search for downstream effectors of ATR-mediated telomerase inhibition at DSBs, related to Figure 5 (A) Western blot for the indicated proteins performed on whole-cell lysates from RPE1-ST cells treated with Cas12a and CRISPR RNAs (crRNAs) targeting RAD51, HSSB1, and RADX or a control (cntrl) crRNA. Ku70, loading control. (B) Quantification of neotelomere formation by TaqMan qPCR in the RPE1-ST cells shown in (A) normalized to cells treated with the control crRNA. (C) Western blot for the indicated proteins performed on whole-cell lysates from RPE1-ST cells treated with Cas12a and crRNAs targeting BLM, WRN, or FANCJ or a control (cntrl) crRNA. γ-tub, loading control. (D) Quantification of neotelomere formation by TaqMan qPCR in the RPE1-ST cells shown in (C) normalized to cells treated with the control crRNA. (E–F) Western blot for the indicated proteins performed on whole-cell lysates from RPE1-ST cells treated with Cas12a and crRNAs targeting XPF (E), SLX4 (F), or a control (cntrl) crRNA. γ-tub, loading control. (G) Quantification of neotelomere formation by TaqMan qPCR in the RPE1-ST cells shown in (E) and (F) normalized to cells treated with the control crRNA. Mean ± SD of at least 3 biological replicates. ns *p* > 0.05, * *p* < 0.05, two-tailed ratio-paired *t*-test in (B), (D), and (G).

## References

1. Ablasser, A., and Chen, Z. J. (2019). cGAS in action: Expanding roles in immunity and inflammation. Science 363, eaat8657.

2. Abreu, E., Aritonovska, E., Reichenbach, P., Cristofari, G., Culp, B., Terns, R. M., Lingner, J., and Terns, M. P. (2010). TIN2-tethered TPP1 recruits human telomerase to telomeres in vivo. Mol Cell Biol 30, 2971–2982.

3. Alexandrov, L. B. et al. (2020). The repertoire of mutational signatures in human cancer. Nature 578, 94–101.

4. Anzalone, A. V. et al. (2019). Search-and-replace genome editing without double-strand breaks or donor DNA. Nature 576, 149–157.

5. Ballif, B. C., Yu, W., Shaw, C. A., Kashork, C. D., and Shaffer, L. G. (2003). Monosomy 1p36 breakpoint junctions suggest pre-meiotic breakage-fusion-bridge cycles are involved in generating terminal deletions. Hum Mol Genet 12, 2153–2165.

6. Bodnar, A. G., Ouellette, M., Frolkis, M., Holt, S. E., Chiu, C. P., Morin, G. B., Harley, C. B., Shay, J. W., Lichtsteiner, S., and Wright, W. E. (1998). Extension of life-span by introduction of telomerase into normal human cells. Science 279, 349–352.

7. Bonaglia, M. C. et al. (2011). Molecular mechanisms generating and stabilizing terminal 22q13 deletions in 44 subjects with Phelan/McDermid syndrome. PLoS Genet 7, e1002173.

8. Boulé, J. B., Vega, L. R., and Zakian, V. A. (2005). The yeast Pif1p helicase removes telomerase from telomeric DNA. Nature 438, 57–61.

9. Boulé, J. B., and Zakian, V. A. (2007). The yeast Pif1p DNA helicase preferentially unwinds RNA DNA substrates. Nucleic Acids Res 35, 5809–5818.

10. Cejka, P., and Symington, L. S. (2021). DNA End Resection: Mechanism and Control. Annu Rev Genet 55, 285–307.

11. Chen, C., and Kolodner, R. D. (1999). Gross chromosomal rearrangements in Saccharomyces cerevisiae replication and recombination defective mutants. Nat Genet 23, 81–85.

12. Chen, L. Y., Redon, S., and Lingner, J. (2012). The human CST complex is a terminator of telomerase activity. Nature 488, 540–544.

13. Chung, W. H., Zhu, Z., Papusha, A., Malkova, A., and Ira, G. (2010). Defective resection at DNA double-strand breaks leads to de novo telomere formation and enhances gene targeting. PLoS Genet 6, e1000948.

14. Cleal, K., Jones, R. E., Grimstead, J. W., Hendrickson, E. A., and Baird, D. M. (2019). Chromothripsis during telomere crisis is independent of NHEJ, and consistent with a replicative origin. Genome Res 29, 737–749.

15. Cristofari, G., and Lingner, J. (2006). Telomere length homeostasis requires that telomerase levels are limiting. EMBO J 25, 565–574.

16. de Lange, T. (2018). Shelterin-Mediated Telomere Protection. Annu Rev Genet 52, 223–247.

17. de Lange, T., and Borst, P. (1982). Genomic environment of the expression-linked extra copies of genes for surface antigens of Trypanosoma brucei resembles the end of a chromosome. Nature 299, 451–453.

18. Dewhurst, S. M., Yao, X., Rosiene, J., Tian, H., Behr, J., Bosco, N., Takai, K. K., de Lange, T., and Imieliński, M. (2021). Structural variant evolution after telomere crisis. Nat Commun 12, 2093.

19. Di Virgilio, M., and Gautier, J. (2005). Repair of double-strand breaks by nonhomologous end joining in the absence of Mre11. J Cell Biol 171, 765–771.

20. Fan, Q., and Yao, M. (1996). New telomere formation coupled with site-specific chromosome breakage in Tetrahymena thermophila. Mol Cell Biol 16, 1267–1274.

21. Fan, Q., and Yao, M. C. (2000). A long stringent sequence signal for programmed chromosome breakage in Tetrahymena thermophila. Nucleic Acids Res 28, 895–900.

22. Feng, J., Funk, W. D., Wang, S. S., Weinrich, S. L., Avilion, A. A., Chiu, C. P., Adams, R. R., Chang, E., Allsopp, R. C., and Yu, J. (1995). The RNA component of human telomerase. Science 269, 1236–1241.

23. Flint, J., Craddock, C. F., Villegas, A., Bentley, D. P., Williams, H. J., Galanello, R., Cao, A., Wood, W. G., Ayyub, H., and Higgs, D. R. (1994). Healing of broken human chromosomes by the addition of telomeric repeats. Am J Hum Genet 55, 505–512.

24. Gerstung, M. et al. (2020). The evolutionary history of 2,658 cancers. Nature 578, 122–128.

25. Ghanim, G. E., Fountain, A. J., van Roon, A. M., Rangan, R., Das, R., Collins, K., and Nguyen, T. H. D. (2021). Structure of human telomerase holoenzyme with bound telomeric DNA. Nature 593, 449–453.

26. Greider, C. W., and Blackburn, E. H. (1985). Identification of a specific telomere terminal transferase activity in Tetrahymena extracts. Cell 43, 405–413.

27. Guilherme, R. S., Hermetz, K. E., Varela, P. T., Perez, A. B., Meloni, V. A., Rudd, M. K., Kulikowski, L. D., and Melaragno, M. I. (2015). Terminal 18q deletions are stabilized by neotelomeres. Mol Cytogenet 8, 32.

28. Hadi, K. et al. (2020). Distinct Classes of Complex Structural Variation Uncovered across Thousands of Cancer Genome Graphs. Cell 183, 197–210.e32.

29. Hahn, W. C., Stewart, S. A., Brooks, M. W., York, S. G., Eaton, E., Kurachi, A., Beijersbergen, R. L., Knoll, J. H., Meyerson, M., and Weinberg, R. A. (1999). Inhibition of telomerase limits the growth of human cancer cells. Nat Med 5, 1164–1170.

30. Hanish, J. P., Yanowitz, J. L., and de Lange, T. (1994). Stringent sequence requirements for the formation of human telomeres. Proc Natl Acad Sci U S A 91, 8861–8865.

31. Hannes, F., Van Houdt, J., Quarrell, O. W., Poot, M., Hochstenbach, R., Fryns, J. P., and Vermeesch, J. R. (2010). Telomere healing following DNA polymerase arrest-induced breakages is likely the main mechanism generating chromosome 4p terminal deletions. Hum Mutat 31, 1343–1351.

32. Hoa, N. N. et al. (2016). Mre11 Is Essential for the Removal of Lethal Topoisomerase 2 Covalent Cleavage Complexes. Mol Cell 64, 580–592.

33. Hockemeyer, D., and Collins, K. (2015). Control of telomerase action at human telomeres. Nat Struct Mol Biol 22, 848–852.

34. Horsley, S. W. et al. (2001). Monosomy for the most telomeric, gene-rich region of the short arm of human chromosome 16 causes minimal phenotypic effects. Eur J Hum Genet 9, 217–225.

35. Jia, P., Chastain, M., Zou, Y., Her, C., and Chai, W. (2017). Human MLH1 suppresses the insertion of telomeric sequences at intra-chromosomal sites in telomerase-expressing cells. Nucleic Acids Res 45, 1219–1232.

36. Kottemann, M. C., and Smogorzewska, A. (2013). Fanconi anaemia and the repair of Watson and Crick DNA crosslinks. Nature 493, 356–363.

37. Lamb, J., Harris, P. C., Wilkie, A. O., Wood, W. G., Dauwerse, J. G., and Higgs, D. R. (1993). De novo truncation of chromosome 16p and healing with (TTAGGG)n in the alpha-thalassemia/mental retardation syndrome (ATR-16). Am J Hum Genet 52, 668–676.

38. Lei, M., Zaug, A. J., Podell, E. R., and Cech, T. R. (2005). Switching human telomerase on and off with hPOT1 protein in vitro. J Biol Chem 280, 20449–20456.

39. Li, T., and Chen, Z. J. (2018). The cGAS-cGAMP-STING pathway connects DNA damage to inflammation, senescence, and cancer. J Exp Med 215, 1287–1299.

40. Liao, S., Tammaro, M., and Yan, H. (2016). The structure of ends determines the pathway choice and Mre11 nuclease dependency of DNA double-strand break repair. Nucleic Acids Res 44, 5689–5701.

41. Lingner, J., Hughes, T. R., Shevchenko, A., Mann, M., Lundblad, V., and Cech, T. R. (1997). Reverse transcriptase motifs in the catalytic subunit of telomerase. Science 276, 561–567.

42. Liu, B., He, Y., Wang, Y., Song, H., Zhou, Z. H., and Feigon, J. (2022). Structure of active human telomerase with telomere shelterin protein TPP1. Nature 604, 578–583.

43. Lydeard, J. R., Lipkin-Moore, Z., Jain, S., Eapen, V. V., and Haber, J. E. (2010). Sgs1 and exo1 redundantly inhibit break-induced replication and de novo telomere addition at broken chromosome ends. PLoS Genet 6, e1000973.

44. MacDougall, C. A., Byun, T. S., Van, C., Yee, M. C., and Cimprich, K. A. (2007). The structural determinants of checkpoint activation. Genes Dev 21, 898–903.

45. Maciejowski, J., and de Lange, T. (2017). Telomeres in cancer: tumour suppression and genome instability. Nat Rev Mol Cell Biol 18, 175–186.

46. Maciejowski, J., Li, Y., Bosco, N., Campbell, P. J., and de Lange, T. (2015). Chromothripsis and Kataegis Induced by Telomere Crisis. Cell 163, 1641–1654.

47. McClintock, B. (1938). The Production of Homozygous Deficient Tissues with Mutant Characteristics by Means of the Aberrant Mitotic Behavior of Ring-Shaped Chromosomes. Genetics 23, 315–376.

48. McClintock, B. (1941). The Stability of Broken Ends of Chromosomes in Zea Mays. Genetics 26, 234–282.

49. McClintock, B. (1983). Letter from Barbara McClintock to Elizabeth H. Blackburn.

50. Mirman, Z., and de Lange, T. (2020). 53BP1: a DSB escort. Genes Dev 34, 7–23.

51. Mirman, Z., Lottersberger, F., Takai, H., Kibe, T., Gong, Y., Takai, K., Bianchi, A., Zimmermann, M., Durocher, D., and de Lange, T. (2018). 53BP1-RIF1-shieldin counteracts DSB resection through CST- and Polα-dependent fill-in. Nature 560, 112–116.

52. Mirman, Z., Sasi, N. K., King, A., Chapman, J. R., and de Lange, T. (2022). 53BP1-shieldin-dependent DSB processing in BRCA1-deficient cells requires CST-Polα-primase fill-in synthesis. Nat Cell Biol 24, 51–61.

53. Morin, G. B. (1991). Recognition of a chromosome truncation site associated with alpha-thalassaemia by human telomerase. Nature 353, 454–456.

54. Motwani, M., Pesiridis, S., and Fitzgerald, K. A. (2019). DNA sensing by the cGAS-STING pathway in health and disease. Nat Rev Genet 20, 657–674.

55. Myung, K., Chen, C., and Kolodner, R. D. (2001). Multiple pathways cooperate in the suppression of genome instability in Saccharomyces cerevisiae. Nature 411, 1073–1076.

56. Nandakumar, J., Bell, C. F., Weidenfeld, I., Zaug, A. J., Leinwand, L. A., and Cech, T. R. (2012). The TEL patch of telomere protein TPP1 mediates telomerase recruitment and processivity. Nature 492, 285–289.

57. Nguyen, T. H. D., Tam, J., Wu, R. A., Greber, B. J., Toso, D., Nogales, E., and Collins, K. (2018). Cryo-EM structure of substrate-bound human telomerase holoenzyme. Nature 557, 190–195.

58. Paiano, J. et al. (2021). Role of 53BP1 in end protection and DNA synthesis at DNA breaks. Genes Dev 35, 1356–1367.

59. Pascolo, E., Wenz, C., Lingner, J., Hauel, N., Priepke, H., Kauffmann, I., Garin-Chesa, P., Rettig, W. J., Damm, K., and Schnapp, A. (2002). Mechanism of human telomerase inhibition by BIBR1532, a synthetic, non-nucleosidic drug candidate. J Biol Chem 277, 15566–15572.

60. Ribeyre, C., and Shore, D. (2013). Regulation of telomere addition at DNA double-strand breaks. Chromosoma 122, 159–173.

61. Rivera, M. A., and Blackburn, E. H. (2004). Processive utilization of the human telomerase template: lack of a requirement for template switching. J Biol Chem 279, 53770–53781.

62. Saldivar, J. C., Cortez, D., and Cimprich, K. A. (2017). The essential kinase ATR: ensuring faithful duplication of a challenging genome. Nat Rev Mol Cell Biol 18, 622–636.

63. Schulz, V. P., and Zakian, V. A. (1994). The saccharomyces PIF1 DNA helicase inhibits telomere elongation and de novo telomere formation. Cell 76, 145–155.

64. Sekne, Z., Ghanim, G. E., van Roon, A. M., and Nguyen, T. H. D. (2022). Structural basis of human telomerase recruitment by TPP1-POT1. Science 375, 1173–1176.

65. Sternberg, S. H., Redding, S., Jinek, M., Greene, E. C., and Doudna, J. A. (2014). DNA interrogation by the CRISPR RNA-guided endonuclease Cas9. Nature 507, 62–67.

66. Stracker, T. H., and Petrini, J. H. (2011). The MRE11 complex: starting from the ends. Nat Rev Mol Cell Biol 12, 90–103.

67. Swarts, D. C., van der Oost, J., and Jinek, M. (2017). Structural Basis for Guide RNA Processing and Seed-Dependent DNA Targeting by CRISPR-Cas12a. Mol Cell 66, 221–233.e4.

68. Varley, H., Di, S., Scherer, S. W., and Royle, N. J. (2000). Characterization of terminal deletions at 7q32 and 22q13.3 healed by De novo telomere addition. Am J Hum Genet 67, 610–622.

69. Wang, F., Stewart, J. A., Kasbek, C., Zhao, Y., Wright, W. E., and Price, C. M. (2012). Human CST has independent functions during telomere duplex replication and C-strand fill-in. Cell Rep 2, 1096–1103.

70. Wilkie, A. O., Lamb, J., Harris, P. C., Finney, R. D., and Higgs, D. R. (1990). A truncated human chromosome 16 associated with alpha thalassaemia is stabilized by addition of telomeric repeat (TTAGGG)n. Nature 346, 868–871.

71. Wong, A. C., Ning, Y., Flint, J., Clark, K., Dumanski, J. P., Ledbetter, D. H., and McDermid, H. E. (1997). Molecular characterization of a 130-kb terminal microdeletion at 22q in a child with mild mental retardation. Am J Hum Genet 60, 113–120.

72. Wong, J. M., and Collins, K. (2006). Telomerase RNA level limits telomere maintenance in X-linked dyskeratosis congenita. Genes Dev 20, 2848–2858.

73. Wu, P., Takai, H., and de Lange, T. (2012). Telomeric 3’ overhangs derive from resection by Exo1 and Apollo and fill-in by POT1b-associated CST. Cell 150, 39–52.

74. Wu, R. A., Upton, H. E., Vogan, J. M., and Collins, K. (2017). Telomerase Mechanism of Telomere Synthesis. Annu Rev Biochem 86, 439–460.

75. Yang, Z., Maciejowski, J., and de Lange, T. (2017). Nuclear Envelope Rupture Is Enhanced by Loss of p53 or Rb. Mol Cancer Res 15, 1579–1586.

76. Yao, M. C., Yao, C. H., and Monks, B. (1990). The controlling sequence for site-specific chromosome breakage in Tetrahymena. Cell 63, 763–772.

77. Yatsenko, S. A., Brundage, E. K., Roney, E. K., Cheung, S. W., Chinault, A. C., and Lupski, J. R. (2009). Molecular mechanisms for subtelomeric rearrangements associated with the 9q34.3 microdeletion syndrome. Hum Mol Genet 18, 1924–1936.

78. Zetsche, B. et al. (2015). Cpf1 is a single RNA-guided endonuclease of a class 2 CRISPR-Cas system. Cell 163, 759–771.

79. Zetsche, B. et al. (2017). Multiplex gene editing by CRISPR-Cpf1 using a single crRNA array. Nat Biotechnol 35, 31–34.

80. Zhang, W., and Durocher, D. (2010). De novo telomere formation is suppressed by the Mec1-dependent inhibition of Cdc13 accumulation at DNA breaks. Genes Dev 24, 502–515.

81. Zhao, Y., Sfeir, A. J., Zou, Y., Buseman, C. M., Chow, T. T., Shay, J. W., and Wright, W. E. (2009). Telomere extension occurs at most chromosome ends and is uncoupled from fill-in in human cancer cells. Cell 138, 463–475.

82. Zhong, F. L., Batista, L. F., Freund, A., Pech, M. F., Venteicher, A. S., and Artandi, S. E. (2012). TPP1 OB-fold domain controls telomere maintenance by recruiting telomerase to chromosome ends. Cell 150, 481–494.

83. Zhou, X. Z., and Lu, K. P. (2001). The Pin2/TRF1-interacting protein PinX1 is a potent telomerase inhibitor. Cell 107, 347–359.

84. Zhu, X., Clarke, R., Puppala, A. K., Chittori, S., Merk, A., Merrill, B. J., Simonović, M., and Subramaniam, S. (2019). Cryo-EM structures reveal coordinated domain motions that govern DNA cleavage by Cas9. Nat Struct Mol Biol 26, 679–685.

